# Robust V(D)J Recombination in the Absence of SA1/2-mediated Chromatin Loops

**DOI:** 10.64898/2026.10.10.757971

**Authors:** Zhuo Yang, Yu Fu, Mariia Mikhova, David Chen, Camille Hebert, Adam Yongxin Ye, Xin Lin, Taimor Williams, Hongli Hu, Zhaoqing Ba, Job Dekker, Frederick W. Alt

## Abstract

The cohesin ring complex organizes and mediates functional activity of the genome through chromatin loop extrusion^1,2^. Cohesin comprises a core SMC1–SMC3–RAD21 ring associated with multiple regulatory proteins, including SA1 and SA2 paralogs that bind mutually-exclusively to a RAD21 site^3,4^. Functional studies of cohesin complexes *in vivo* often employ inactivation of particular components^2,3^. However, inactivation of SA1 or SA2 in mice is limited by mouse embryonic lethality^5–7^ and, for dual SA1/2-inactivation by cycling cell lethality^8–10^. For these reasons, it remains unknown whether the requirement for SA1/2 in loop extrusion *in vitro*^11,12^ applies *in vivo*^1,2^, and whether these factors have overlapping or distinct functions in regulating this process^3^. Here, we employ G0/G1-arrested *v-Abl* pro-B lines, which are viable in the absence of cohesin-complex factors^13,14^, to dissect SA1 and SA2 functions in V(D)J recombination, a strictly G0/G1-specific process that harnesses loop extrusion to effect distinct *Igh* and *Igκ* locus recombination mechanisms^15,16^. Robust long-range *Igh* or *Igκ* V(D)J recombination continued unabated in the absence of SA1 or SA2. However, inactivation of both abrogated most positioned chromatin loops genome-wide and also abolished long-range *Igh* V(D)J recombination with a phenotype similar to RAD21 (cohesin) inactivation^13^. Yet, robust *Igκ* V(D)J recombination continued upon dual SA1/2-inactivation, which along with Hi-C findings, demonstrates ongoing loop-extrusion in the absence of SA1/2. Correspondingly, replacement of weak *Igh* V(D)J recombination signal sequences (RSSs) that target V(D)J recombination activity, with strong *Igκ* counterparts^15^ restored long-range *Igh* V(D)J recombination in SA1/2 inactivated cells. We conclude that SA1/2 together maintain loop extrusion impediments and genome-wide architecture in G0/G1-arrested cells; and, in their absence, loop extrusion proceeds unimpeded. Remarkably, strong RSSs allow *Igκ* to function normally in the absence of internal positioned loop structure across the 3.1-mega-base Vκ locus. We discuss implications of our findings for long-range gene regulation in G0/G1-arrested cells more broadly.

## Background

Cycling cell studies implicated distinct functions for SA1 or SA2 cohesin complexes. SA1 complexes have been implicated in preferentially supporting stable CTCF structural factor-associated architecture and larger, more stable loops such as convergent CTCF binding element (CBE)-based loop domains^3,10,17,18^. SA2 has been implicated in contributing to smaller, more dynamic sub-loops, such as those involved in enhancer-promoter interactions and context-dependent gene regulation^3,19–21^. Elegant single-molecule *in vitro* studies demonstrated that either an SA1- or SA2-containing cohesin complex is essential for cohesin-mediated loop extrusion of DNA^11,12,22^. Despite these advances, little is known about how these paralogs act individually or redundantly over long range in given genomic loci to control structural and functional outcomes in physiological processes. Also, the functional requirement for SA1 or SA2 for cohesin-mediated chromatin loop extrusion in living cells remains speculative^1,2^. Moreover, despite most cells in the mammalian body being viably arrested in G0/G1^23^, few studies examined functions of cohesin complex factors in regulating loop extrusion in G0/G1 cells. In this regard, cycling cells express both extrusive cohesin that carries out loop extrusion and cohesive cohesin involved in sister chromatid cohesion and is only present in cycling cells^24^, which could potentially confound some cohesin-complex functions in cycling cell studies.

V(D)J recombination, which occurs strictly in the G0/G1 cell cycle phase^25^, can be activated in G0/G1-arrested Abelson murine leukemia virus-transformed pro-B cell lines (“*v-Abl* cells”)^26^. G0/G1-arrested *v-Abl* cells undergo cohesin loop extrusion-mediated V(D)J recombination orchestrated by RAG-endonuclease bound to recombination centers (RCs) formed over J_H_s and Jκ clusters, respectively, at the downstream ends of the multi-megabase (Mb) *Igh* and, for primary rearrangements, *Igκ* loci^15,16,27,28^ (Extended Data Fig. 1a,b). RAG cleavage activity is targeted by recombination signal sequences (RSSs) that have two distinct forms denoted 12RSSs and 23RSSs; RAG binds and cleaves only bound pairs of 12 and 23 RSSs^29–31^. 100-plus V_H_s, 10 D_H_s, and 4 J_H_s are linearly organized across a 2.7 Mb upstream *Igh* region (Extended Data Fig. 1a, panel (i)). V_H_s and J_H_s have 23RSSs and D_H_s have 12RSSs on both sides with the downstream 12RSS in convergent orientation with J_H_ 23RSSs and upstream 12RSS in convergent orientation with V_H_ downstream RSSs, guiding strictly deletional D_H_-to-J_H_ and then V_H_-to-DJ_H_ joining to generate a V(D)J exon (Extended Data Fig. 1a, panel (ii)). *Igκ* V(D)J recombination employs 100-plus Vκs flanked by 12RSSs and 4 functional downstream Jκs flanked by 23RSSs spread over 3 Mb locus (Extended Data Fig. 1b, panel (i)). Most proximal and distal Vκs are oriented for deletional joining to Jκs; while most middle Vκs are orientated for inversional joining (Extended Data Fig. 1b, panel (ii)). *Igh* V_H_, D_H_, and J_H_ segments have relatively weak RSSs and rely on CBE/ transcription-based loop extrusion impediments to increase interaction with the RAG-bound J_H_-based RC during one cohesin-loop-based linear-RAG scanning-mediated rearrangement^13,14,27^. Dozens of proximal V_H_s have RSS-associated CBEs^27^, that impede upstream RAG scanning until these and other impediments are neutralized in normal pro-B cells, by developmental WAPL cohesin unloader down-regulation^14,32^, or in *v-Abl* lines by ectopic WAPL depletion^14^. High level expression of the WAPL in *v-Abl* cells limits cohesin-mediated loop extrusion-based RAG scanning past scanning impediments^14^, including the IGCR1 CBE-based RAG scanning impediment that separates the downstream D_H_ and J_H_ containing domain from the upstream V_H_ domain and restricts robust RAG J_H_-based RC scanning to D_H_ segments (Extended Data Fig. 1a, panel (i))^27,33^. Primary Vκ to Jκ rearrangement involves a two cohesin loop-based rearrangement process in which Vκs, oriented in both deletional and inversional orientations access the primary Jκ-based RC by diffusion across the *Cer/Sis* CBE-based diffusion platform just upstream of the Jκ RC (Extended Data Fig. 1b, panel (i))^15^. Such two-loop diffusion-based access for Vκs across the 3.1 Mb Vκ locus is possible for primary Vκ-to-Jκ V(D)J recombination, but not for that of *Igh*, because Vκ and Jκ RSSs are immensely more robust than those of V_H_s, D_H_s, and J_H_s^15^.

### Effect of SA1 and/or SA2 inactivation on *Igh* and *Igκ* internal interaction domains

While inactivation of RAD21 is lethal in cycling cells, G0/G1-arrested *v-Abl* cells in which RAD21^13^, or several other cohesin-complex factors^13,14^, were individually inactivated remain viable, allowing V(D)J recombination functions to be studied. Thus *v-Abl* cells should offer an ideal system for studying SA1 and SA2 functions in the context of V(D)J recombination in viably G0/G1-arrested cells that do not undergo the rapid lethality associated with dual SA1/2 inactivation that has limited analyses in cycling cells. We employed G0/G1-arrested 129SV mouse genetic background *v-Abl* cells to assess effects of inactivating SA1 or SA2, individually or in pairs, on V(D)J recombination across the *Igh* and *Igκ* loci, a process in which cohesin-mediated loop extrusion plays distinct, fundamental mechanistic roles^13–16,34^. We generated degron-dTAG-based conditional protein-depletion “SA1-degron” and “SA2-degron”, gene-knockout “*SA1^-/-^*” and “*SA2^-/-^*” deficient *v-Abl* cells, as well as “*SA1*^-/-^/SA2-degron” or “*SA2*^-/-^/SA1-degron” Rag endonuclease-deficient *v-Abl* pro-B cells (Extended Data Fig. 2a-c). RAG-deficient *v-Abl* cell lines are used for all experiments described in this study, except HTGTS-V(D)J-Seq experiments in which RAG expression is ectopically introduced, to prevent confounding effects of endogenous V(D)J rearrangements in the *Igh* and *Igκ* loci^27,34,35^ (See Methods, for details). Effective elimination of SA1 and/or SA2 in “*SA1^-/-^*” and “*SA2^-/-^*” *v-Abl* cells, as well as in “SA1-degron”, “SA2-degron”, “*SA1*^-/-^/SA2-degron”, and “*SA2*^-/-^/SA1-degron” cells following 6 hours of dTAG-13, was confirmed by western blotting (Extended Data Fig. 2d) and by absence of chromatin-bound SA1 and SA2 across the *Igh* and *Igκ* loci evidenced by ChIP-seq (Extended Data Fig. 2g-h). Elimination of SA1, SA2 or both paralogs had minimal impact on viability of the G0/G1-arrested *v-Abl* cells (Extended Data Fig. 2e-f). All experiments discussed below were done in the G0/G1-arresested *v-Abl* cells described above and all experiments described for *SA1*^-/-^/SA2-degron, or *SA2*^-/-^/SA1-degron cells will be referred to as “dual” inactivation. Following G0/G1 arrest, we employed Hi-C^36,37^ to examine *Igh* or *Igκ* loop structure in single SA1-depleted or SA2-depleted *v-Abl* cells and in dual SA1/2 inactivated *SA1*^-/-^/SA2-depleted *v-Abl* cells.

*Igh* locus Hi-C analyses confirmed that parental cells exhibited distinct internal interaction domains and RC-associated chromatin structures spanning regions both upstream and downstream of the RC^32,38^ (Fig. 1a,b; panel (i)). Upon individual depletion of SA1 or SA2 in SA1-depleted or SA2-depleted *v-Abl* lines, *Igh* internal loop structures were largely maintained (Fig. 1a,b; panels (ii-iii)). Following SA1/2 dual inactivation in *SA1*^-/-^/SA2-depleted *v-Abl* lines, the majority of these local *Igh* chromatin structures were abrogated, indicating a broad loss of chromatin loop-extrusion impediments (Fig. 1a,b; panel (iv)). Similarly, in the *Igκ* locus, parental cells displayed prominent internal contact domains, consistent with previous observations in primary pre-B cells, where such structures have been proposed to juxtapose Vκ and Jκ segments to facilitate efficient Vκ-to-Jκ joining^38^ (Fig. 1c,d; panel (i)). Upon individual depletion of SA1 or SA2 in SA1-degron or SA2-degron *v-Abl* cells, *Igκ* internal loop structures were largely maintained, other than the appearance of a modest *Igκ* sub-TAD located within distal and middle Vκ clusters following SA2 depletion (Fig. 1c,d; panels (i-iii)). Upon SA1/2 dual inactivation in *SA1*^-/-^/SA2-degron *v-Abl* cells, all *Igκ*-internal positioned loop structures were largely eliminated (Fig. 1c,d; panel (iv)); however, concomitant with the loss of *Igκ* internal positioned loops, a large TAD that spanning the Cer/Sis region to the distal Vκ region^15,39^, that is barely visible in parental and single depleted cells, became prominent (Fig. 1c,d; panel (iv)). As compared to the Hi-C contact map of *SA1*^-/-^/SA2-degron dual inactivated *v-Abl* cells, the disappearance of this prominent TAD in RAD21-depleted cells was the only clear difference (Fig. 1c,d; compared panels iv to v). Overall, these studies indicate that, following SA1/2 dual inactivation, the majority of local *Igh* and *Igκ* positioned loop structures were abrogated, indicating a broad loss of chromatin loop-extrusion impediments across the *Igh* and *Igκ* locus (Fig. 1). Further, the large TAD in the *Igκ* locus detected following SA1/2 dual inactivation is a structure that requires RAD21. Overall interactions across *Igh* and *Igκ* were higher after SA1/2 dual inactivation as compared to Rad21 depletion (Extended Data Fig. 3), pointing to remaining cohesin-dependent contacts throughout both loci after combined loss of SA1 and SA2.

**Fig. 1.**
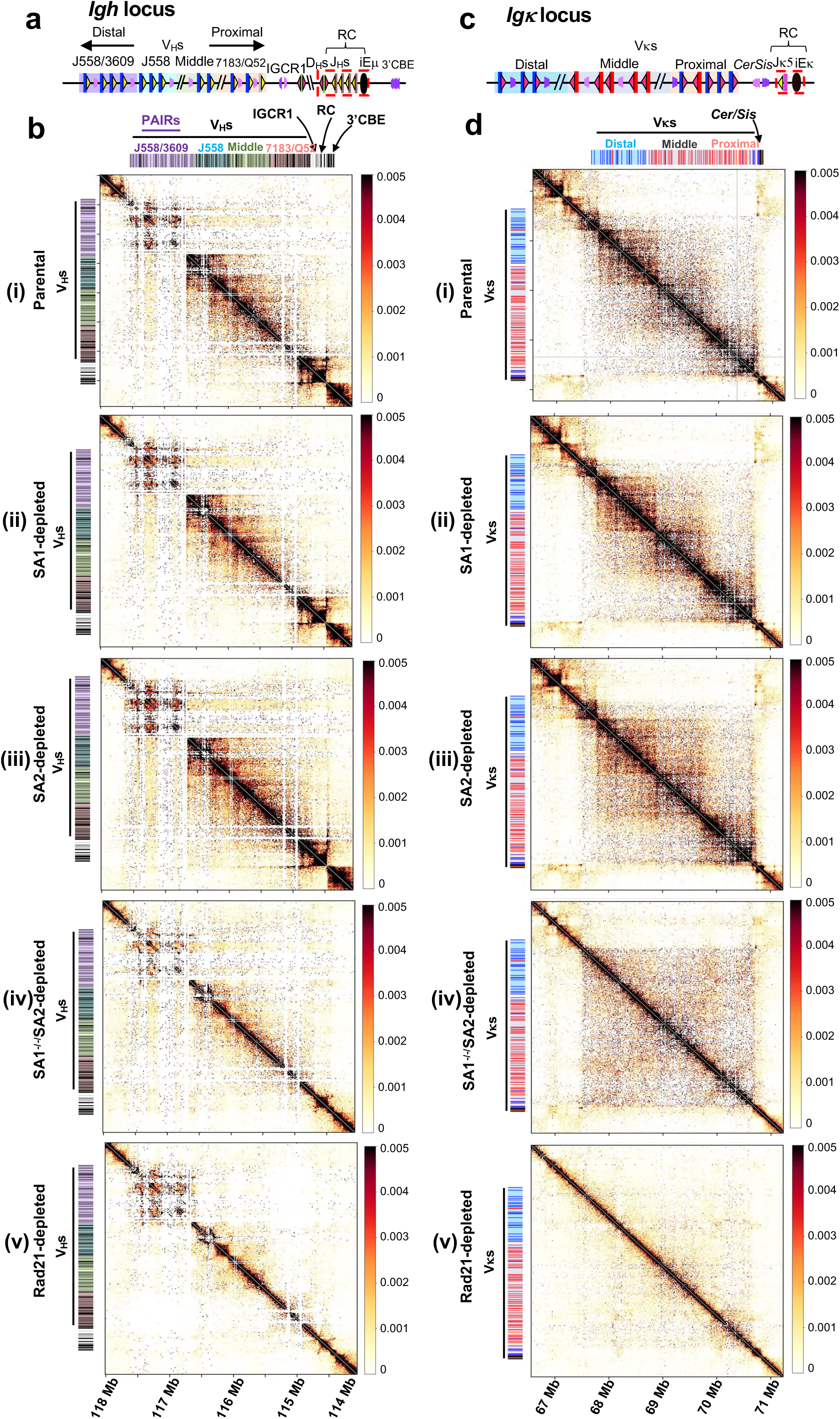
SA1/2 inactivation neutralizes most loop-extrusion impediments in *Igh* and *Igκ* loci. **a**, Schematic of the mouse *Igh* locus (not to scale) showing V_H_s, D_H_s, J_H_s, with the *Igh* RC that comprises the J_H_-proximal D_H_, four J_H_s, and the intronic enhancer (iEμ) highlighted in dashed red box. The V_H_s are divided into four V_H_ domains: proximal 7183/Q52 (pink shadow), middle (green shadow), distal J558 and distal J558/3609 (blue and purple shadow). The light and dark purple arrowheads indicate organization of *Igh* CBEs including two divergently oriented CBEs within the *Igh* IGCR1. The yellow and red triangles represent the position and orientation of bona fide 23RSSs and 12RSSs, respectively. **b**, Hi-C contact maps of the *Igh* (**a**) loci at 10 kb resolution in (i) parental cells, (ii) SA1-depleted cells, (iii) SA2-depleted cells, (iv) *SA1^-/-^*/SA2-depleted cells, and (v) RAD21-depleted cells. The SA1-degron, SA2-degron, and *SA1^-/-^*/SA2-degron cells were treated with dTAG-13 before G0/G1 arrest, respectively. RAD21-degron cells were treated with IAA before G0/G1 arrest^13^. See text for more detailed description. **c**, Schematic of the mouse *Igκ* locus, showing the relative positions of Vκs: proximal (pink shadow), middle (gray shadow), and distal (blue shadow). The single Jκ5 and the *Igκ* intronic enhancer (iEκ) contribute to the *Igκ* RC (dashed red box). *Cer* and *Sis* elements, located between the Vκs and the *Igκ* RC, each contain two CBEs (light and dark purple arrowheads) that are convergently oriented. Vκ 12RSS (red triangles), Jκ 23RSS (yellow triangles), and other Vκ locus CBEs (purple arrowheads) are indicated. **d**, Hi-C contact maps of the *Igκ* loci. Locus orientations and annotations are shown above the maps. Contact frequencies were plotted from genome-wide ICE-normalized matrices using a linear color scale from 0 to 0.005. See text for more detailed description.

To further clarify the effects of SA1/2 dual inactivation on the *Igh* and *Igκ*, we used high-resolution 3C-HTGTS^13–15,27^ to examine organizational features at high resolution. For *Igh*, we used CBE1 of the CBE-based IGCR1 element^13,14,27^ and the iEμ enhancer^13,14,27^ of the *Igh* RC as baits (Extended Data Fig. 4a). Consistent with loss of major positioned loop structures observed by Hi-C, SA1/2 inactivation in the *SA1*^-/-^/SA2-degron line abolished IGCR1 CBE1-mediated interactions with proximal V_H_s, the RC, and the 3′CBEs, along with eliminating low-level enhancer-mediated transcription-associated interactions within the downstream region (Extended Data Fig. 4b, panels (i-iv)). Moreover, SA1/2 inactivation in *SA1*^-/-^/SA2-degron line markedly reduced interactions of the iEμ/RC with proximal V_H_s, IGCR1 and downstream 3′CBEs (Extended Data Fig. 4b, panels (v-viii)). For *Igκ*, we used the Cer CBE1^15,16^, a middle Vκ locus CBE, and the RC iEκ enhancer^15,16^ as 3C-HTGTS baits (Extended Data Fig. 4c). Dual inactivation of SA1/2 in the *SA1*^-/-^/SA2-degron *v-Abl* cells nearly eliminated Cer CBE1 interactions with CBE-associated sites (Extended Data Fig. 4d, panels (i-iv)); notably, most remaining sites contain either E2A, transcription, both, or a combination of the three (Supplementary information Fig.1). Consistent with Hi-C data, the middle Vκ CBE bait revealed a range of interaction sites in the distal/middle Vκ regions, substantially contributed by CBE-based interactions, that largely disappeared upon dual SA1/2 inactivation in the *SA1*^-/-^/SA2-degron *v-Abl* cells (Extended Data Fig. 4d, panels (v-viii); and Fig. 1d). The iEκ/RC bait, as expected^15,16^, did not interact with upstream Vκs^15,16^ (Extended Data Fig. 4d, panel (ix)). However, this bait strongly interacted with downstream regulatory elements, including 3′Eκ and Ed, as well as with downstream CBEs (Extended Data Fig. 4d, panel (x)). Notably, after SA1/2 dual inactivation in *SA1*^-/-^/SA2-degron *v-Abl* line, downstream CBE interactions were lost; whereas interactions with 3′Eκ and Ed were still detected (Extended Data Fig. 4d, panel (xii)). 3C-HTGTS analyses in RAD21-depleted *v-Abl* cells employing a Cer-CBE1 bait confirmed Hi-C findings, showing RAD21 inactivation, in contrast to SA1/2 inactivation, abrogated Cer-CBE1 interactions across the Vκ locus (Supplementary information Fig. 2a,b). Thus, in contrast to SA1/2 inactivation, RAD21-inactivation abrogates all long-range interactions. Overall, our Hi-C and 3C-HTGTS findings indicate that SA1/2 inactivation neutralizes many loop-extrusion impediments across *Igh* and *Igκ* loci; but that some elements, including *Cer/Sis* and RCs retain impediment activity. Additional findings, including functional data that support this conclusion is presented below.

**Fig. 2.**
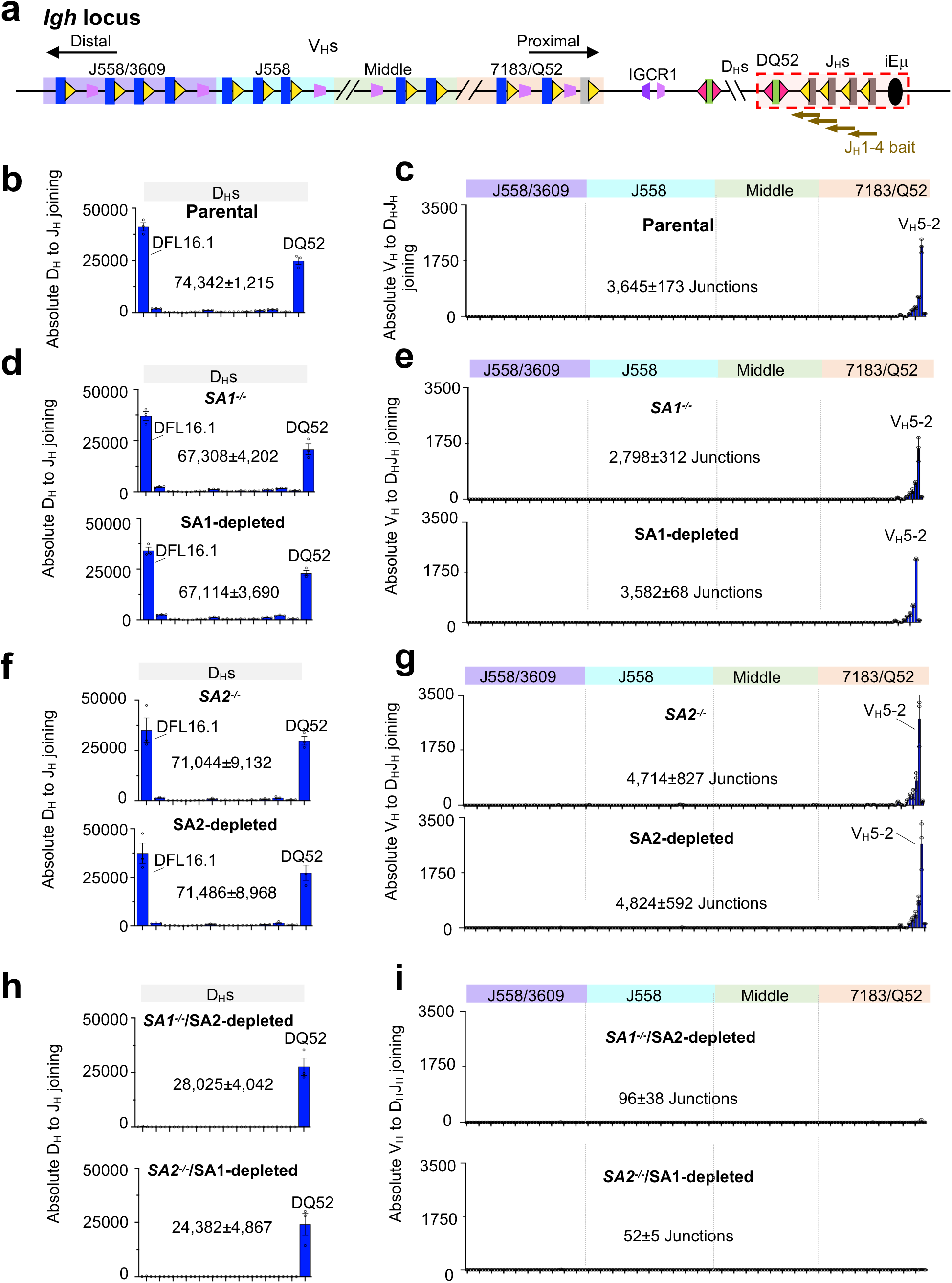
Effects of SA1 and/or SA2 inactivation on *Igh* V(D)J recombination. **a**, Schematic of the mouse *Igh* locus. Locations of selected D_H_, V_H_, and J_H_ segments are indicated. Brown arrows under J_H_s coding end bait primers used to generate HTGTS-V(D)J-seq libraries. Other details in legend to Figure 1a. **b-i**, Absolute utilization of all D_H_s in D_H_-to-J_H_ rearrangements (**b, d, f, h**) and V_H_s in V_H_-to-DJ_H_ rearrangements (**c, e, g, i**) in different *v-Abl* lines indicated as: parental (**b, c**); *SA1^-/-^*(**d**, **e**; upper panels) and SA1-depleted (**d, e**; lower panel); *SA2^-/-^* (**f**, **g**; upper panels) and SA2-depleted (**f, g**; lower panel); *SA1^-/-^*/SA2-depleted (**h**, **i**; upper panels) and *SA2^-/-^*/SA1-depleted (**h**, **i**; lower panels). In the *Igh* locus, essentially all rearrangements are deletional and indicated in blue bar. Total junction numbers are indicated in each panel. All D_H_ and V_H_ usage patterns in different single mutants or depletions are highly similar to those of parental cells, as assessed by two-sided Pearson correlation. For D_H_ usage pattern compared with parental: r = 1 for *SA1^-/-^*; r = 1.00 for SA1-depleted; r = 0.99 for *SA2^-/-^*; and r = 0.99 for SA2-depleted cells, with P < 0.001 for all comparisons. For V_H_ usage pattern compared with parental: r = 1.00 for all indicated single mutant or depletion conditions, with P < 0.001. Moreover, D_H_ and J_H_ rearrangement patterns were same between *SA1^-/-^*/SA2-depleted and *SA2^-/-^*/SA1-depleted cells (r = 1.00, P < 0.001). In contrast, SA1/2 dual inactivation substantially altered both D_H_ and V_H_ rearrangement pattern relative to parental cells, with r = 0.46 for D_H_ usage, and r = 0.50 for V_H_ usage, both with P < 0.001. DQ52 usage was not significantly changed after SA1/2 dual inactivation compared with parental cells: 1.11-fold change for *SA1^-/-^*/SA2-depleted cells (P = 0.55) and 0.97-fold change for *SA2^-/-^*/SA1-depleted cells (P = 0.90). Data are shown as mean ± s.e.m. from three biological replicates. Fold-change P values were calculated using unpaired, two-tailed Welch’s t-tests.

### Effects of SA1 and/or SA2 inactivation on *Igh* V(D)J recombination

In *v-Abl* cells RAG predominantly utilizes DQ52, which lies in the RC and gains RAG access by short-range diffusion; and distal DFL16.1, which gains RAG access during scanning at its location adjacent to the IGCR1 loop extrusion impediment that blocks upstream scanning^13,26,33^ (Fig. 2a,b). In *v-Abl* cells, the proximal V_H_5-2 (V_H_81X) is used at low levels via “leaky” RAG scanning through IGCR1, with utilization dependent upon its RSS-associated CBE^27^ (Fig. 2a,c). Several proximal V_H_s just upstream of V_H_5-2 are utilized at even lower levels and no further upstream V_H_s are utilized in *v-Abl* cells, except after down-regulation of loop extrusion impediments by ectopic inactivation of WAPL^14^ or CTCF^13^. In G0/G1-arrested *v-Abl* cells, these *Igh* parental line D_H_-to-J_H_ and V_H_-to-DJ_H_ recombination patterns and levels were fully maintained upon SA1 or SA2 inactivation in SA1-degron and *SA1^-/-^ v-Abl* cells, as well as in SA2-degron and *SA2^-/-^ v-Abl* cells (Fig. 2a-g). Thus, both SA1 and SA2, individually, can maintain normal *Igh* V(D)J recombination in G0/G1-arrested 129SV *v-Abl* cells and are functionally redundant in this role. In striking contrast, SA1/2 dual inactivation in *SA1*^-/-^/SA2-degron or *SA2*^-/-^/SA1-degron *v-Abl* cells abrogated both DFL16.1-to-J_H_ and all V_H_-to-DJ_H_ rearrangements (Fig. 2h,i). However, DQ52 rearrangement by diffusional access was still utilized for D_H_-to-J_H_ joining in SA1/2 dual inactivated cells (Fig. 2h), indicating that *Igh* RC activity is maintained in the dual absence of SA1 and SA2. We note that the *Igh* locus phenotype of dual SA1 and SA2 inactivation is essentially the same as that observed for RAD21 inactivation^13^.

### Effects of SA1 and/or SA2 inactivation on *Igκ* primary V(D)J recombination

We studied primary Vκ to Jκ joining in a single Jκ5-containing *v-Abl* line that can only undergo primary rearrangements^15^. Primary *Igκ* rearrangements involve joining of deletional-oriented and inversional-oriented Vκs across the locus by short-range diffusion-based access to the RAG-bound Jκ5-based RC as they are extruded past the Cer/Sis diffusion platform^15^ (Fig. 3a,b). In contrast to *Igh*, such Vκ to Jκ joining occurs in the presence of high WAPL levels, due at least in part to weaker CBEs across the Vκ locus and much stronger *Igκ* RSSs^15^. We found no difference in Vκ-to-Jκ primary rearrangement patterns upon SA1 or SA2 inactivation in SA1-depleted cells and *SA1^-/-^ v-Abl* cells (Fig. 3c), as well as in SA2-depleted cells and *SA2^-/-^ v-Abl* cells (Fig. 3d), as compared to those of parental cells with the only difference being a modest increase in levels of Vκ rearrangements in SA2-depleted cells and *SA2^-/-^ v-Abl* cells (Fig. 3d; and Supplementary information Fig. 3a-c, panels i-ii). The slightly increased Vκ-Jκ joining levels with no change in Vκ-Jκ joining pattern across the locus is most consistent with a slight increase in *Igκ* RC activity^14^. Remarkably, despite absence of internal Vκ locus positioned loop structures in SA1/2 dual inactivated *SA1*^-/-^/SA2-degron or *SA2*^-/-^/SA1-degron *v-Abl* cells (Fig. 1d, panel (iv)), we found relatively normal levels of primary Vκ-to-Jκ deletional and inversional rearrangements across the *Igκ* locus, but with a shift towards more distal Vκ utilization (Fig. 3e and Supplementary information Fig. 3d, panels i-ii). In contrast to SA1/2 inactivation, RAD21 inactivation abrogated *Igκ* RC transcription and Vκ to Jκ rearrangement (Supplementary information Fig. 2c,d; see legend for further discussion). Together, the above findings indicate that loop extrusion-mediated RAG scanning and diffusion-based deletional and inversional Vκ capture mediated by the *Cer/Sis* diffusion platform are largely maintained in SA1/2 dual inactivated cells. The finding that primary deletional and inversional Vκ-to-Jκ rearrangement persists in SA1/2 dual inactivated cells is consistent with the strong *Cer/Sis* CBEs and *Igκ* RC remaining functional in the absence of both paralogs. Notably, these findings also clearly demonstrate that chromatin loop extrusion continues to function in the absence of SA1 and SA2 *in vivo* in G0/G1-arrested *v-Abl* cells. Moreover, the *Igh* and *Igκ* dual-inactivation findings, considered together, strongly indicate that dual absence of SA1 and SA2 leads to accelerated loop extrusion through the two loci due to neutralization of most loop extrusion scanning impediments.

**Fig. 3.**
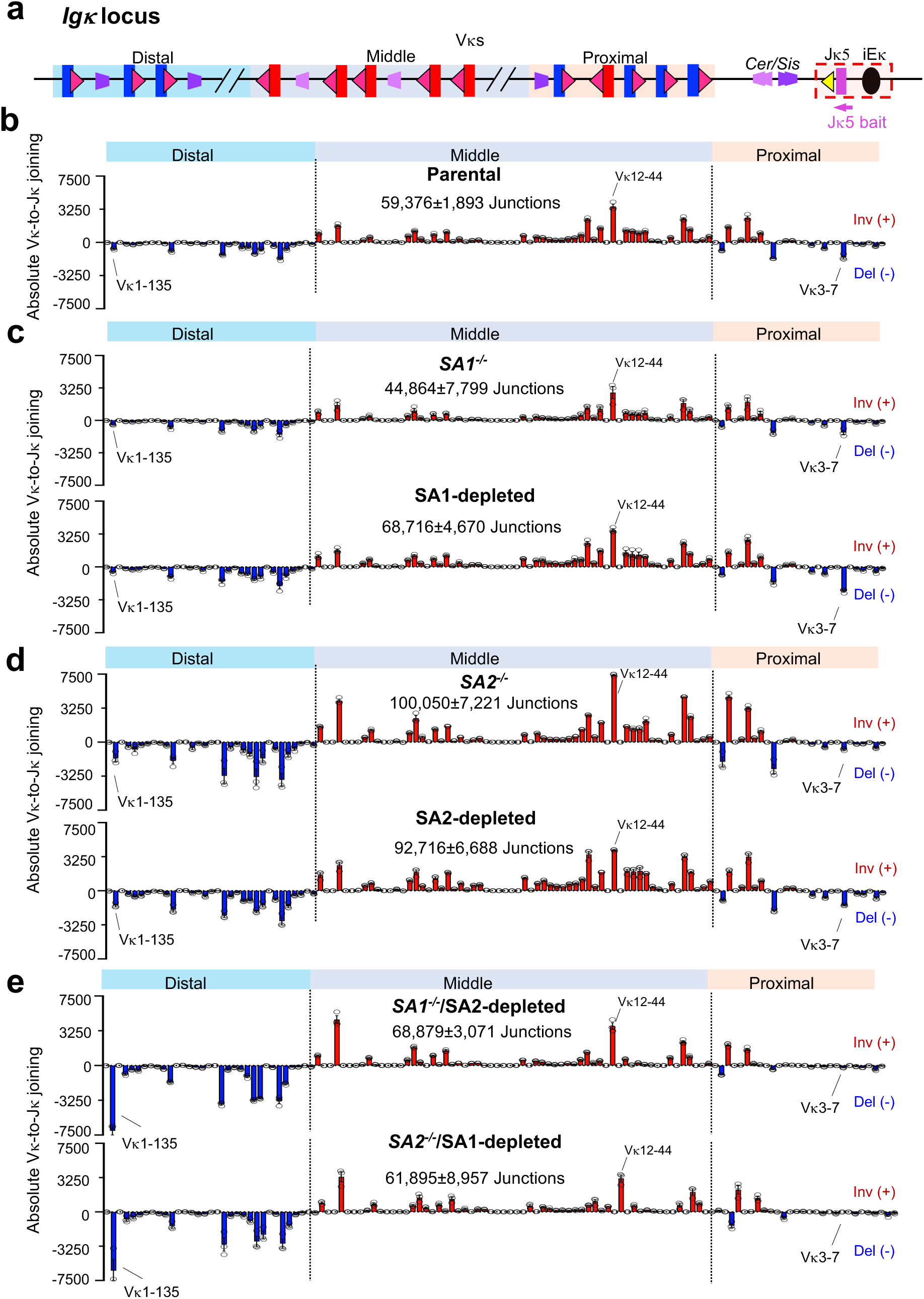
Effects of SA1 and/or SA2 inactivation on *Igκ* V(D)J recombination. **a**, Schematic of the mouse *Igκ* locus in a single Jκ5 *v-Abl* cell line. Light purple arrow under Jκ5 coding end (CE) bait primer used to generate HTGTS-V(D)J-seq libraries. Other details in legend to Figure 1c. **b-e**, Absolute utilization of all Vκs in *Igκ* locus in different *v-Abl* cells as indicated: parental (**b**); *SA1^-/-^* (**c**; upper panels) and SA1-depleted (**c**; lower panel); *SA2^-/-^* (**d**; upper panels) and SA2-depleted (**d**; lower panel); *SA1^-/-^*/SA2-depleted (**e**; upper panels) and *SA2^-/-^*/SA1-depleted (**e**; lower panels). In all panels, deletional Vκ-Jκ joins are plotted below the x-axis in blue and labeled “Del (−)”, inversional Vκ-Jκ joins are plotted above the x-axis in red and labeled “Inv (+)”. Total junction numbers are shown in each panel. Vκ rearrangement patterns in different single mutants or depletions are highly similar to those of parental cells, as assessed by two-sided Pearson correlation. Compared to parental: r = 0.99 for *SA1^-/-^*, r = 0.99 for SA1-depleted, r = 0.96 for *SA2^-/-^*, and r = 0.98 for SA2-depleted cells, with P < 0.001 for all comparisons. The Vκ rearrangement patterns were same between *SA1^-/-^*/SA2-depleted and *SA2^-/-^*/SA1-depleted lines (r = 0.99, P < 0.001). By contrast, SA1/2 dual inactivation resulted in a distinct Vκ rearrangement pattern relative to parental cells (r = 0.75, P < 0.001). Total Vκ-Jκ junction numbers were comparable among parental cells, *SA1^-/-^* and SA1-depleted lines, and SA1/2 dual-inactivation lines, see Supplementary Table 2. The exceptions were *SA2^-/-^*and SA2-depleted lines, which showed significant increases from parental cells (*SA2^-/-^*: 1.7-fold increase, P = 0.02; SA2-depleted: 1.6-fold increase, P = 0.03, respectively). Data are presented as mean ± s.e.m. P values were calculated using an unpaired, two-tailed Welch’s t-test (n = 3).

### Impact of leaky depletion of SA1 in *SA2^-/-^* cells or SA2 in *SA1^-/-^* cells

Western blot analysis revealed low-level leaky depletion without dTAG-13 treatment in the single SA1-degron and SA2-degron cells, as well as in the untreated *SA1*^-/-^/SA2-degron or *SA2*^-/-^/SA1-degron cells (Extended Data Fig. 2d). Therefore, we examined whether such leaky depletion affected *Igh* or *Igκ* V(D)J recombination by comparing levels and patterns in G0/G1-arrested untreated single SA1-degron or SA2-degron cells lines, as well as in untreated *SA1*^-/-^/SA2-degron cells and *SA2*^-/-^/SA1-degron cells. Untreated single SA1-degron or SA2-degron cells had *Igh* and *Igκ* rearrangement patterns that were very similar to parental cells (Extended Data Fig. 5a; Extended Data Fig. 6a), as expected since complete depletion or inactivation of one or the other by targeted deletion has no phenotype (Fig. 2b-g and Fig. 3b-d). In contrast, untreated *SA1*^-/-^/SA2-degron and *SA2*^-/-^/SA1-degron cells showed, nearly identical impacts on both loci as compared to parental or untreated single-degron levels (Extended Data Fig. 5a-b; Extended Data Fig. 6a-b). In *Igh,* the ratio of DFL16.1 to DQ52 DJ_H_ rearrangements was reversed, compared to those of parental cells, in both leaky genotypes, with DFL16.1 levels decreased and those of DQ52 increased, a phenotype consistent with weakening of IGCR1 CBEs^14^ (Extended Data Fig. 5c). However, the level of V(D)J rearrangements was not significantly different from those of parental, single-degron untreated cells, or *SA1*^-/-^/SA2-degron and *SA2*^-/-^/SA1-degron untreated cells (Extended Data Fig. 5a, panels (ii,vi,vi); Extended Data Fig. 5b, panels (ii,vi)), consistent with the overall level of DJ_H_ rearrangement not significantly changing (Extended Data Fig. 5b, panels (i,iii)). In the *Igκ*, overall Vκ rearrangement levels increased over two-fold in both *SA1*^-/-^/SA2-degron and *SA2*^-/-^/SA1-degron genotypes, although rearrangement patterns were similar to parental cells in both cases (Extended Data Fig. 6a-c). The parallel *Igh* phenotypes of the *SA1*^-/-^/SA2-degron and *SA2^-/-^*/SA1-degron cells indicate that one overlapping function of the two paralogs involves maintenance of CBE impediment activity. The significant increase in *Igκ* recombination levels, without a change in Vκ rearrangement patterns, for both *SA1*^-/-^/SA2-degron and *SA2^-/-^*/SA1-degron *v-Abl* cells, as mentioned for complete SA2 inactivation above (Fig. 3d), is most consistent with a change in RC activity; but further studies are needed to elucidate the mechanism.

### Impact of SA2 depletion in C57BL/6-background *v-Abl* cells

A very recent study published shortly before paper was submitted reported that single SA2 inactivation in C57BL/6-background G0/G1-arrested *v-Abl* cells expanded low level V_H_ utilization beyond proximal V_H_s^40^. In contrast, our analyses in 129SV-background *v-Abl* cells showed that single SA1 or SA2 inactivation had no significant effect on the dominant utilization of the very most proximal V_H_s (Fig. 2a-g). Previous studies suggested somewhat diminished utilization of V_H_5-2 compared to immediately upstream proximal V_H_s in C57BL/6-background versus 129SV-background *v-Abl* cells^13^. To test whether genetic background differences might contribute to differential effects of SA2 inactivation on V_H_ utilization in *v-Abl* cells, we generated SA1-degron and SA2-degron C57BL/6-background *v-Abl* cell lines and assessed *Igh* V(D)J rearrangement patterns in parental, SA1-inativated, or SA2-inactivated cells after G0/G1 arrested. Overall V_H_-to-DJ_H_ recombination patterns were highly similar between parental, SA1-depleted, and SA2-depleted C57BL/6-background *v-Abl* cells (Extended Data Fig. 7a-c, panels (i-ii)). Nevertheless, SA2-depleted, but not SA1-depleted cells showed low, but detectable, extension of V_H_-to-DJ_H_ recombination beyond the proximal V_H_ region (Extended Data Fig. 7c, panels (i-ii)). Due to the low level of these extended V_H_ rearrangements in C57BL/6 SA2-inactivated cells, we also show these patterns on a broken scale, (Extended Data Fig. 7a-c, panel (iii)) as was done in the prior study^40^ to highlight the low, but significant extension of V_H_ utilization in the SA2-inactivated C57BL/6 background *v-Abl* cells (see legend of Extended Data Fig. 7 for details). For comparison with the C57BL/6 background *v-Abl* cell findings, we have also similarly presented the parental, SA1-depleted, and SA2-depleted 129SV *v-Abl* cell HTGTS-V(D)-Seq data from Fig. 2 (Extended Data Fig. 7d-f, panels (i-iii)). Consistent with previous observations of differences between C57BL/6- and 129SV-background *v-Abl* cells^13,14^, parental C57BL/6-background *v-Abl* cells showed significantly increased relative utilization of proximal V_H_s just upstream of V_H_5-2 as compared to parental 129SV-background *v-Abl* cells (Extended Data Fig. 7a,d, compare panel (i); see legend for details). However, in contrast to the effect of SA2 inactivation observed in C57BL/6-background cells, SA2-depleted 129SV-background *v-Abl* cells showed no significant difference in upstream V_H_ usage compared with parental or SA1-depleted 129SV-background cells (Extended Data Fig. 7d-f, panel (iii)). Together, the 129SV background versus C57BL/6 background studies confirm a subtle difference in the effects of SA2 versus SA1 inactivation during *Igh* V(D)J recombination in the C57BL/6 background *v-Abl* cells.

### Linear scanning-mediated secondary *Igκ* rearrangements in SA1/2-inactivated cells

In normal *v-Abl* cells or bone marrow pre-B cells, deletion of *Cer/Sis* leads to robust rearrangement of proximal Vκs via linear RAG scanning into the proximal Vκ locus; utilization is limited to the very most proximal Vκs due to rapid saturation of V(D)J recombination events by the strong Vκ and Jκ RSSs during direct one-loop-based linear scanning^16^. This rearrangement pattern mimics that of secondary *Igκ* rearrangements across the Vκ locus^16^. In this context *Cer/Sis* deletion by primary deletional rearrangements or displacement by primary inversional rearrangements serves as a developmental switch to convert the *Igκ* locus from using a two-loop domain primary rearrangement mechanism into a locus that uses a one-loop-based linear scanning mechanism for secondary rearrangements^16^. To test the effects of *Cer/Sis* deletion in the context of SA1/2 dual inactivation, we generated *SA1*^-/-^/SA2-degron, *Cer/Sis*-deleted *v-Abl* cells. Strikingly, *SA1*^-/-^/SA2-inactivated, *Cer/Sis*-deleted *v-Abl* cells similarly promoted robust rearrangement of the very most proximal Vκs to the “ectopic secondary” Jκ-RC^16^, as parental *Cer/Sis*-deleted *v-Abl* cells, albeit with rearrangements extended slightly more distally for the dual SA1/2 inactivated cells (Fig. 4a,b). Overall, our findings on *Igκ* V(D)J recombination in the absence of SA1/2, indicate that both primary and secondary Vκ-to-Jκ recombination mechanisms function in the absence of most internal Vκ positioned loop structure.

**Fig. 4.**
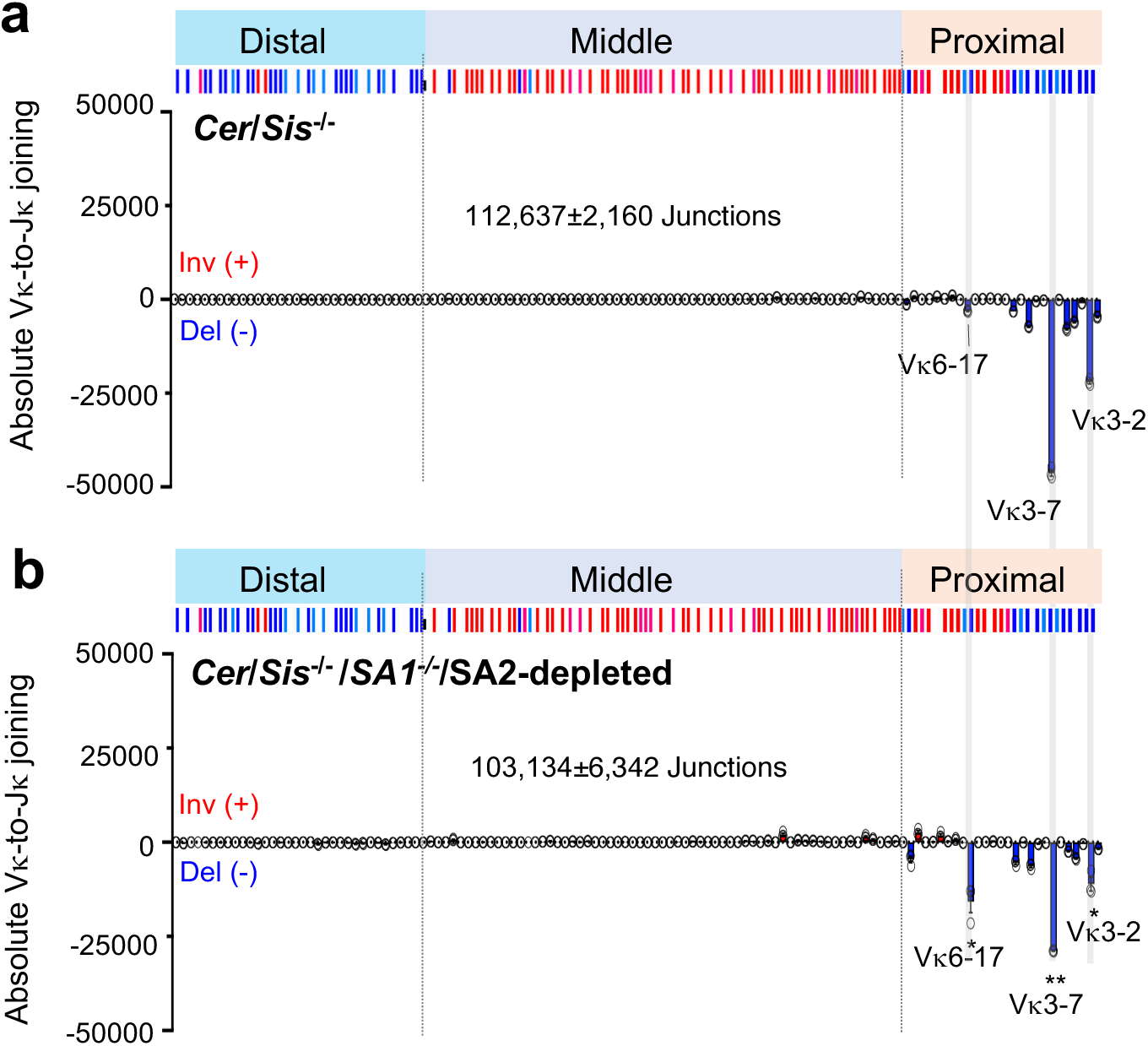
Robust deletional proximal Vκs rearrangement upon *Cer*/*Sis* knockout in SA1/2 inactivation cells. **a**,**b**, Absolute utilization of Vκs in *Cer*/*Sis*^-/-^ (**a**) or *Cer*/*Sis*^-/-^/*SA1*^-/-^/SA2-depleted (**b**) cell lines. Blue tracks below the x-axis indicate deletional Vκ-Jκ joins “Del (−)”, and red tracks above the x-axis indicate inversional Vκ-Jκ joins “Inv (+)” in all panels. Total junction numbers are shown in each panel. Locations of selected Vκ segments are indicated. Vκ rearrangement patterns are highly similar between *Cer*/*Sis*^-/-^ (**a**) and *Cer*/*Sis*^-/-^/*SA1*^-/-^/SA2-depleted cells (**b**) (two-sided Pearson’s r = 0.90, P < 0.001), although rearrangements extended modestly more distally after SA1/2 dual inactivation. This extension was associated with modestly reduced usage of proximal Vκ3-2 and Vκ3-7 segments (1.99-fold, P = 0.02; and 1.60-fold, P = 0.0022, respectively) and increased usage of the more distal Vκ6-17 (5.13-fold, P = 0.046). Vκ junction utilization data are presented as mean ± s.e.m. from 3 biological replicates. P values were calculated using an unpaired, two-tailed Welch’s t-test (n = 3) (*P ≤ 0.05, **P ≤ 0.01, ***P ≤ 0.001).

### Strong *Igκ* RSS promote *Igh* V(D)J rearrangement upon SA1/2 inactivation

To determine whether strong RSSs can drive V(D)J recombination in the *Igh* via a one-loop-based linear RAG-scanning mechanism, we generated a pre-rearranged DQ52-J_H_4 “DJ_H_” allele in the *SA1*^-/-^/SA2-degron *v-Abl* cell line, referred to as the “DJ_H_ *SA1*^-/-^/SA2-degron” line (Fig. 5a). In the DJ_H_ *SA1*^-/-^/SA2-degron *v-Abl* untreated line, proximal impediments, particularly the V_H_5-2 RSS-associated CBE, restrict RAG scanning to the very proximal V_H_s, resulting in almost exclusively proximal V_H_5-2 rearrangements via “leaky” RAG scanning through IGCR1^15^ (Fig. 5b). SA1/2 dual inactivation in the DJ_H_ *SA1*^-/-^/SA2-degron *v-Abl* line abrogated proximal V_H_ rearrangements (Fig. 5c). To test influence of strong *Igκ* RSSs, we modified the parental DJ_H_ *SA1*^-/-^/SA2-degron line by replacing the weak DQ52 upstream RSS^15^ with a strong Vκ12-44-12RSS to create the “VκRSS-DJ_H_ *SA1*^-/-^/SA2-degron” *v-Abl* line (Fig. 5d). As anticipated, this line, due to the now strong DJ_H_RC RSS, had an approximately 8-fold increased V_H_5-2 utilization (Fig. 5e) which was nearly abrogated upon dual SA1/2 inactivation (Fig. 5f). We then replaced the RSS of the V_H_8-12, located 2 Mb upstream of the *Igh* RC, with a strong Jκ1-23 RSS to generate the Jκ/VκRSS-DJ_H_ *SA1*^-/-^/SA2-degron line (Fig. 5g). In the Jκ/VκRSS-DJ_H_*SA1*^-/-^/SA2-degron untreated line, the rearrangement pattern was quite similar to that of the VκRSS-DJ_H_ *SA1*^-/-^/SA2-degron untreated line, with very little upstream V_H_8-12 rearrangement despite the strong Jκ1-23RSS (Fig. 5h). However, upon SA1/2 inactivation in the Jκ/VκRSS-DJ_H_ *SA1*^-/-^/SA2-degron line, V_H_5-2 rearrangements decreased to very low levels, while V_H_-to-DJ_H_ recombination of the distal V_H_8-12 with the strong Jκ1-23RSS was robustly increased (Fig. 5i). We note that the 3C-HTGTS profile, when baited from the *Igh* RC (iEμ), revealed that the strong *Igκ* RSSs did not promote V_H_8-12 interaction with the *Igh* RC (Extended Data Fig. 8a), consistent with our prior findings that robust *Vκ* RSSs do not require impediment activity to increase their utilization during linear-scanning mediated secondary *Igκ* rearrangements^15^.

**Fig. 5.**
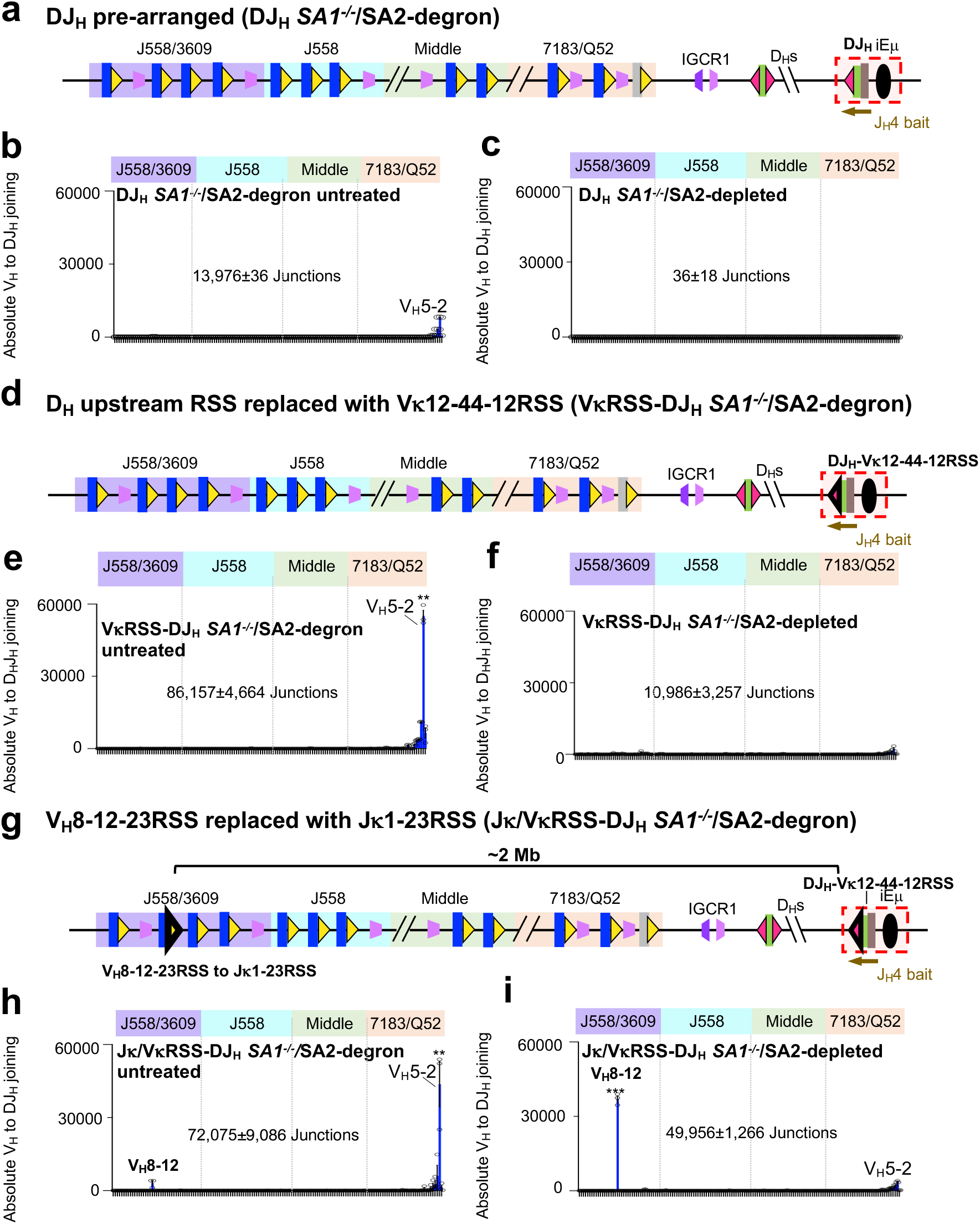
Strong *Igκ* RSS promotes long-range V_H_ usage upon SA1/2 dual inactivation. **a**, Diagram of DQ52-J_H_4 pre-rearranged allele in *SA1*^-/-^/SA2-degron *v-Abl* cell line referred to as the “DJ_H_ *SA1*^-/-^/SA2-degron”. The DJ_H_ RC is indicated in red dashed box. Brown arrow under J_H_4 coding end (CE) indicates bait primer used to generate HTGTS-V(D)J-seq libraries. **b, c**, Absolute utilization of V_H_s in *Igh* in *SA1*^-/-^/SA2-degron untreated cell line (**b**) and in *SA1*^-/-^/SA2-depleted cell line (**c**). **d**, Diagram showing single RSS replacement in DJ_H_ *SA1*^-/-^/SA2-degron *v-Abl* lines, referred to as “VκRSS-DJ_H_ *SA1*^-/-^/SA2-degron”. The upstream RSS of DQ52 was replaced with the Vκ12-44-12RSS, indicated by a bold-black-outlined red triangle. **e, f**, Absolute utilization of V_H_s in *Igh* of VκRSS-DJ_H_ *SA1*^-/-^/SA2-degron untreated (**e**) and VκRSS-DJ_H_ *SA1*^-/-^/SA2-depleted cells (**f**). After single RSS replacement, total V_H_s junction numbers in VκRSS-DJ_H_ *SA1*^-/-^/SA2-degron untreated cells increased significantly (6.16-fold increase, P = 0.0041) compared with DJ_H_ *SA1*^-/-^/SA2-degron untreated cells. The usage of V_H_5-2 increased 6.74-fold (P = 0.0024) compared with its usage in DJ_H_ *SA1*^-/-^/SA2-degron untreated cells. **g**, Diagram illustrating the double RSS replacement strategy used to generate the Jκ/VκRSS-DJ_H_ *SA1*^-/-^/SA2-degron *v-Abl* cell line. Junction numbers are shown in each panel. Locations of selected V_H_ segments are indicated. The V_H_8-12-23RSS was replaced with the Jκ1-23RSS in the VκRSS-DJ_H_ *SA1*^-/-^/SA2-degron cell line, indicated by a bold-black-outlined yellow triangle. **h, i**, Absolute utilization of V_H_s in Jκ/VκRSS-DJ_H_ *SA1*^-/-^/SA2-degron *v-Abl* cell untreated (**h**) and Jκ/VκRSS-DJ_H_ *SA1*^-/-^/SA2-depleted cells (**i**). The usage of V_H_8-12 in Jκ/VκRSS-DJ_H_ *SA1*^-/-^/SA2-depelted cell line increased more than 1100-fold (P = 0.0012) compared with DJ_H_ *SA1*^-/-^/SA2-degron untreated cells. Utilization of V_H_ data are presented as mean ± s.e.m. from 3 biological replicates. P values were calculated using an unpaired, two-tailed Welch’s t-test (n = 3) (*P ≤ 0.05, **P ≤ 0.01, ***P ≤ 0.001).

To confirm dependence of this long-range V_H_-to-DJ_H_ recombination on loop extrusion, we generated a *v-Abl* line that allowed RAD21 inactivation in the Jκ/VκRSS-DJ_H_ *SA1*^-/-^/SA2-degron line, which we refer to as the Jκ/VκRSS-DJ_H_ *SA1*^-/-^/SA2-degron/RAD21-degron line (Extended Data Fig. 8b-e). As expected RAD21 depletion abolished all V_H_ rearrangements in this cell line (Extended Data Fig. 8g). To confirm RC activity, we further deleted the 100 kb interval between V_H_5-2 and the DJ_H_ RC (Extended Data Fig. 8f). In this position, V_H_5-2 rearrangement was robust following cohesin depletion (Extended Data Fig. 8f,h), consistent with short-range diffusion-based access of V_H_5-2 to the active DJ_H_ RC^27^. Together, these results demonstrate that SA1/2 dual inactivation neutralizes RAG-scanning impediments across the *Igh* locus, including IGCR1 and proximal V_H_ RSS-associated CBEs, allowing unimpeded RAG scanning over the 2 Mb intervening region between the DJ_H_ RC and V_H_8-12, during which strong *Igκ* RSSs can mediate robust V_H_8-12-to-DJ_H_ joining. Thus, *Igκ* RSS replacements allow the *Igh* to undergo robust long-range V(D)J recombination in the absence SA1 and SA2, and, correspondingly, in the absence of internal positioned loop domains formed by stalling of cohesin at impediments.

## Discussion

Consistent with our Hi-C studies that indicate SA1 and SA2 are functionally redundant in maintaining most *Igh* and *Igκ* internal stalled loop structures. We found no appreciable effect of the individual inactivation of SA1 or of SA2 on *Igh* or *Igκ* V(D)J recombination in G0/G1-arrested 129SV *v-Abl* cells (Fig. 2a-g; and Fig. 3a-d). We did confirm a subtle impact of SA2 versus SA1 inactivation on V(D)J recombination in C57BL/6 *v-Abl* cells, potentially reflecting differences in proximal RAG scanning impediment strength in the two genetic backgrounds (Extended Data Fig. 7). However, consistent with most local *Igh* positioned loop structures disappearing in the absence of both SA1 and SA2, RAG scanning-based *Igh* V(D)J recombination was abrogated in the absence of both factors (Fig. 2i). Notably, the *Igh* V(D)J recombination phenotype in the dual absence of SA1 and SA2 was identical to that of RAD21 inactivation^13^, which at first glance may suggest absence of loop extrusion. Yet, Vκ-to-Jκ two-loop-based primary and one-loop-based secondary-like V(D)J recombination mechanisms^15,16^ persisted (Fig. 3; and Fig. 4), robustly, in the absence of SA1 and SA2, despite absence of internal positioned loop structures proposed to be key for such processes^38^. Remarkably, in dual SA1/2 inactivated G0/G1-arrested 129SV *v-Abl* cells, *Igκ* RSS replacements in *Igh* allowed unimpeded loop-extrusion-mediated RAG-scanning over a 2 Mb intervening V_H_ locus region to promote robust rearrangement of a distal V_H_ to a DJ_H_RC (Fig. 5i). Considered together, *Igh* and *Igκ* locus dual-inactivation findings indicate that absence of both SA1 and SA2 leads to accelerated cohesin-mediated loop extrusion through these two loci due to neutralization of most CBE-based and transcription-based loop extrusion scanning impediments (Extended data Fig.9a,b). In this context, powerful *Igκ* RSSs and the still functional *Cer/Sis* diffusion platform and *Igκ* RC, maintain both robust Vκ to Jκ primary and secondary joining (Extended data Fig.9b). In contrast, linear RAG *Igh* scanning was abrogated by dual SA1/2 inactivation due to the elimination of scanning impediments required to promote weak *Igh* RSS interaction with the still active *Igh* RC during linear RAG scanning (Extended Data Fig. 9a, panel (i)), unless rescued by complementary *Igκ* RSS replacements (Extended Data Fig. 9a, panel (ii)). Our studies demonstrate that, in contrast to *in vitro* findings^11,12,22^, cohesin-mediated loop extrusion remains active in the absence of SA1 and SA2 in viably G0/G1-arrested *v-Abl* cells. Moreover, it is striking that strong functional elements, such as *Igκ* RSSs provide robust functional interactions during unimpeded long-range cohesin-mediated loop extrusion, a regulatory strategy that theoretically may have been adopted by other loci. Finally, it is notable that impediment activity of certain key extrusion impediments in the *Igh* and *Igκ* loci functionally persists in the dual absence of SA1 and SA2, including the CBE-based *Cer/Sis* diffusion platform and the transcription-based *Igh* and *Igκ* RCs. Further genome-wide analyses of SA1/2-inactivated, G0/G1-arrested *v-Abl* lines, and other G0/G1-arrested cell types^41^, may elucidate whether other types of key functional elements remain active in other loci in the combined absence of both SA1 and SA2 cohesin complexes.

## Supporting information

Supplementary Information.zip

## Acknowledgements

We thank members of the Alt laboratory for contributions to the study, particularly Chengbin Jing for sharing the dTAG-degron backbone plasmids, Chengxian Xu for providing the total thymocytes cells for western blotting. We thank Cornelis Murre for kindly sharing the Hi-C protocol. We thank Yu Zhang for sharing *SA2^-/-^* data from C57BL/6-background *v-Abl* cells before publication. This work was supported by NIH Grant R01 AI020047 (to F.W.A) and R01 HG003143 (to J.D.). J.D. and F.W.A. are Investigators of the Howard Hughes Medical Institute.

## Author contributions

F.W.A. and Z.Y. designed the study. Z.Y. performed experiments with assistance from M.M., D.C., and C.H. H.H. performed the HTGTS and 3C-HTGTS of RAD21 inactivation. Z.B. performed the Gro-seq of RAD21 inactivation. X.L., T.W., Y.F., and J.D. applied bioinformatics pipelines for data analysis. A.Y.Y. designed bioinformatics pipelines and did statistical analyses. Z.Y., Y.F., J.D., and F.W.A. analyzed and interpreted data and designed figures. F.W.A. and Z.Y. drafted the paper, and Y.F. and J.D. further polished the paper.

## Additional information

Supplementary information is available for this paper.

Correspondence and requests for materials should be addressed to F.W.A.

## Methods

### Experimental procedures

Sample size was not predetermined using statistical methods. No randomization was performed, and investigators were not blinded to group allocation or outcome assessment.

### Generation of SA1 or/and SA2 single and double inactivation *v-Abl* pro-B lines and related derivatives

We used a previously established *Rag2^-/-^*; *Eμ-Bcl^2+^*129SV background *v-Abl* pro-B cell line derived from the 129SV background as the parental line^13,15^. For the single SA1 or SA2 inactivation cell line, we performed two strategies: 1, dTAG degron system to deplete SA1 or SA2 protein in *v-Abl* cell. Targeting constructs were generated to introduce FKBP12^F36V^ sequences into the N terminus of the endogenous mouse SA1 or SA2 locus. The Donor plasmids with mCherry or blasticidin resistance (BSD) cassettes were assembled as illustrated in Extended Data Fig. 2b. Parental *Rag2^-/-^*; *Eμ-Bcl2^+^*cells were nucleofected with SA1-degron-mCherry and SA1-degron-BSD or SA2-degron-mCherry and SA2-degron-BSD donor plasmids, together with the corresponding Cas9/sgRNA vectors. Transfected cells were selected with 16.7 μg/ml blasticidin (Invitrogen, no. R21001) for 3 days, followed by fluorescence-activated cell sorting (FACS) to isolate mCherry-positive cells. Single-cell clones were obtained by limiting dilution, and positive clones were screened by PCR genotyping, verified by Sanger sequencing, and by western blotting. The resulting single SA1 or SA2 inactivation cell lines were referred to as SA1-degron or SA2-degron and used for subsequent experiments. The same strategy was used for generation of SA1-degron or SA2-degron line in *Rag1^-/-^*; *Eμ-Bcl2^+^* C57BL/6 background *v-Abl* cells. 2, CRISPR Cas9/sgRNA targeting to knock out a critical function domain in SA1 or SA2 gene. Two Cas9/sgRNA were designed in the upstream and the downstream of stromalin-conserved domain (SCD), as shown in Extended Data Fig. 2c, which is required for cohesin-SA complex function.

To generate SA1/2 dual-inactivation lines, SA1 or SA2 gene was further knocked out in the SA1-degron or SA2-degron background using Cas9/sgRNAs, and referred to as the *SA1^-/-^*/SA2-degron or *SA2^-/-^*/SA1-degron lines. The *SA1^-/-^*/SA2-degron and *SA2^-/-^*/SA1-degron cell lines with dTAG-13 treatment were named SA1/2 dual inactivated cells. RAD21-degron targeting constructs were generated similarly, except that the FKBP12^F36V^ sequence was inserted at the C terminus of endogenous RAD21, followed by a GFP cassette for screening. A similar strategy was used for generation of *SA1^-/-^*/SA2-degron in *Cer/Sis* deletion^15^ *v-Abl* cells. The IGCR1 deletion cell line was generated as described^13^. All sgRNAs and oligonucleotides sequences used in this study are listed in Supplementary Table 1. *TdT* gene expression and protein levels were detected in this 129SV background *v-Abl* cells, data shown in Supplementary information Figure 4.

### Generation of DJ_H_ pre-rearranged *v-Abl* cell lines and its derivatives

DJ_H_ pre-rearranged *v-Abl* cell lines were generated from *SA1^-/-^*/SA2-degron *v-Abl* cells as described^15,27^. RSS replacement modifications were generated by Cas9/sgRNA-mediated genome editing using ssODNs as donor templates, as described previously^15^. Specifically, the DQ52 upstream RSS was replaced with Vκ12-44-12RSS to generate the VκRSS-DJ_H_ *SA1^-/-^*/SA2-degron line, and the V_H_8-12-23RSS was replaced with Jκ1-12RSS to generate the Jκ/VκRSS-DJ_H_ *SA1^-/-^*/SA2-degron line. The sequences of all sgRNAs and oligonucleotides used in this study are provided in Supplementary Table 1.

### dTAG-mediated single, double, and triple inactivation of SA1, SA2, and RAD21

To induce degradation of FKBP-tagged SA1 and/or SA2, SA1-degron, SA2-degron and SA1/2-inactivated cells were treated with dTAG-13 (Sigma-Aldrich, SML2601) at a final concentration of 0.5 μM. For triple degradation of RAD21, SA1, and SA2, cells were treated with both dTAG-13 and dTAGv (Tocris, 6914), each at a final concentration of 0.5 μM. Stocks of dTAG-13 and dTAGv were prepared at 1,000 × concentration in DMSO. For G0/G1-phase arrest, cells were treated with 3 μM STI-571 for 4 days, while dTAG compounds were maintained in the culture medium. At the end of the treatment period, cells were collected for flow-cytometric and immunoblot analyses to confirm G0/G1 arrest and protein depletion before downstream assays.

### Cell viability measurement

The parental, SA1-degron, SA2-degron, *SA1^-/-^, SA2^-/-^, SA1^-/-^*/SA2-degron, *SA2^-/-^*/SA1-degron, and Jκ/VκRSS-DJ_H_ *SA1^-/-^*/SA2-depleted/RAD21-depleted triple inactivated cells were first incubated with dTAG-13 and/or dTAGv for 6 hours, after which STI-571 treatment was initiated and designated as the 0-hour timepoint. Cell viability over the subsequent 4 days was assessed by FACS as described previously^13^.

### Western blotting

For immunoblotting, 1 × 10^6^ *v-Abl* cells (DMSO or dTAG-13 and/or dTAGv-treated for 6 h) were harvested and lysed as previously described^13,14^. Primary antibodies used were anti-SA1 (Abcam, ab4457), anti-SA2 (Cell Signaling Technology, no. 5882S), anti-RAD21 (Abcam, ab992), TdT antibody (BioLegend, #659002), Wapl antibody (Thermo Fisher Scientific, #PA5-38024), and anti-β-actin (Cell Signaling Technology, no. 5125S). HRP-conjugated secondary antibodies included anti-rabbit HRP (Cell Signaling Technology, no. 7074S) and anti-goat HRP (Thermo Fisher Scientific, no. A15999).

### HTGTS-V(D)J-seq and data analyses

HTGTS-V(D)J-seq libraries were generated as previously described^15,16,42,43^. For RAG2-deficient 129SV background *v-Abl* cell lines, RAG2 was reconstituted by retroviral infection^13,15^. For RAG1-deficient C57BL/6 *v-Abl* cell lines, RAG1 and RAG2 were reconstituted via retroviral infection of cells^14^. For rearrangement analyses, data from 129SV background *v-Abl* cells were aligned to the AJ851868/mm9 hybrid (mm9_AJ851868ins) genome, and data from C57BL/6 *v-Abl* cells were aligned to the mm9 genome. For other *Igh* rearrangement analyses in the DJ_H_ *v-Abl* cell line, reads were aligned to the mm9_AJ851868ins.J_H_4_DQ52_pre_arranged genome, including all duplicate junctions as previously described^43^ (for updated pipeline considerations see: https://github.com/Yyx2626/HTGTS_related), while analyses in VκRSS_DJ_H_, Jκ/VκRSS_DJ_H_, and Trunc/ Jκ/VκRSS_DJ_H_ *SA1^-/-^*/SA2-degron/RAD21-degron *v-Abl* lines were aligned to the mm9_AJ851868ins.J_H_4_DQ52_pre_arranged.Vκ12-44RSS, mm9_AJ851868ins.J_H_4_DQ52_pre_arranged.Vκ12-44RSS.V_H_8-12RSS-Jκ1RSS, and mm9_AJ851868ins.J_H_4_DQ52_pre_arranged.Vκ12-44RSS.V_H_8-12_Jκ1RSS.tunc genomes, respectively. To quantify absolute levels of V(D)J recombination, every HTGTS-V(D)J-seq library was normalized to 280 k reads. Junctions were assigned as deletional or inversional according to the relative orientation of the prey RSS and the bait RSS, as described previously^15,16^.

Relative V(D)J usage patterns were quantified following a previously published HTGTS-V(D)J-seq analysis strategy^14,15,44^. Briefly, 20 k junctions within each library were used to calculate the fractional contribution of each V_H_, D_H_, and Vκ segment, allowing comparison of relative usage patterns independently of total recombination levels.

### 3C-HTGTS and data analyses

3C-HTGTS was performed as previously described^27^ using G0/G1-arrested, RAG2-deficient *v-Abl* cells^27^. Reference genomes were the same as those used for HTGTS-V(D)J-seq analyses. The high peaks in 3C-HTGTS libraries upstream of the bait site were removed^15,16^. Specifically, peaks were removed from the following regions: chr6:70674900-70675400 for iEκ-baited libraries; chr6:70658556-70659968 for Cer CBE1-baited libraries; chr6:70665114-70665735 for Sis CBE2-baited libraries; chr12:114669228-114670236 for iEμ-baited libraries, and chr12:114740073-114741599 for IGCR1 CBE1-baited libraries. Only junctions located within the genomic regions encompassing the full Ig loci (chr6:64,515,000-73,877,000 for *Igκ* and chr12:111,453,935-120,640,000 for *Igh*) were retained for analysis (see details in Code Availability). Processed 3C-HTGTS libraries were then normalized to the total junction count of the smallest library within each comparison set.

### ChIP-seq and data analyses

For ChIP-seq, 20 million G0/G1-arrested parental or SA1/2-inactivated cells (*SA1^-/-^*/SA2-degron line) with dTAG-13 treatment, were collected and processed according to a previously described protocol^13,14^. The antibodies against RAD21 (ab992, Abcam), CTCF (07-729, Millipore), anti-SA1 (Abcam, ab4457), and anti-SA2 (Cell Signaling Technology, no. 5882S) were used to detect the corresponding proteins. ChIP-seq libraries were sequenced on an Illumina NextSeq550 or Illumina NextSeq2000 platform with 150-bp paired-end reads.

### GRO-seq and data analyses

For GRO-seq, 10 million of the G0/G1-arrested parental cells or SA1/2-inactivated cells (*SA1^-/-^*/SA2-degron cells) with dTAG-13 treatment, were collected and processed as previously described^14,34,45^. The resultant GRO-seq libraries were sequenced by a paired-end 150-bp sequencing kit on Illumina NextSeq550 or Illumina NextSeq2000 platform. RAD21-depleted GRO-seq data were downloaded from our previously published dataset^13^.

### Hi-C

Two biological replicates of 10 million G0/G1-arrested cells from each indicated condition were harvested, crosslinked, and processed for Hi-C using the Arima-HiC Kit according to the manufacturer’s instructions^46^. Crosslinked cells were processed for Hi-C using the Arima-HiC Kit (A510008) according to the manufacturer’s instructions. For library preparation, 3.5 µg of proximally ligated DNA was sonicated using a Diagenode Bioruptor to an average fragment size of ∼400 bp (30 s on/ 60 s off for 5 cycles at low power), followed by size selection with AMPure XP beads (Beckman Coulter, A63881). A total of 1.4 µg of fragmented DNA was used for library construction, as amounts closer to 2 µg typically yield higher-complexity Hi-C libraries. Libraries were prepared using the NEBNext Ultra II DNA Library Prep Kit (NEB, E7645) following the Arima-HiC protocol, and sequencing was performed with 150-bp paired-end reads (PE150) on an Illumina NextSeq2000 platform.

### Hi-C data processing

Hi-C data was processed with the distiller-nf (v0.3.4)^47^ pipeline: paired-end 150 bp reads were mapped to the reference genome mm9_AJ851868ins using bwa-mem^48^. Then alignments were pairtools^49^ parsed into interactions with the additional ‘-walk-policy all’ option to keep pairwise and multiway interactions. After deduplication, unique pairs with MAPQ>30 were kept and aggregated into binned contact matrices in the mcool^50^ format at different resolutions. Contact matrices were iteratively balanced (ICEed)^51^ at all resolutions with the default parameters. If multiple Hi-C libraries were shown in one panel, the same number of total valid pairs were randomly sampled from each library at the specified resolution. Chromosome X and Y were excluded in all downstream analyses. For Hi-C data Scaling: Contact frequency (P) as a function of genomic distance (s) and the derivative of the P (s) curve were calculated at 1 kb resolution with cooltools^52^ ‘expected_cis’ function and smoothed in log space using only normalized cis interactions within each chromosome or each specified region.

### Quantification and statistical analysis

For HTGTS, 3C-HTGTS, ChIP-seq, and GRO-seq analyses, each plotting region was divided into either 1,000 or 200 bins using R version 3.6.1, as indicated, or visualized with error bars using a modified version of Integrative Genomics Viewer (IGV: https://github.com/Yyx2626/igv). Signals were first normalized within each replicate and then averaged across three biological replicates. Data are shown as mean ± s.e.m. The V(D)J rearrangement graphs were generated using GraphPad Prism 10. The rearrangement levels were compared between samples using unpaired two-sided Welch’s t-tests, with P values listed in the relevant figure legends. Similarity of V_H_ or Vκ usage patterns was quantified by Pearson correlation after averaging across replicates, and the corresponding r and P values are reported in the figure legends.

## Data availability

High-throughput sequencing data reported in this study have been deposited in the Gene Expression Omnibus (GEO). The associated accession numbers are GSE344621 for HTGTS-V(D)J-seq, GSE343143 for 3C-HTGTS, GSE343144 for ChIP-seq, GSE342264 for GRO-seq, and GSE343090 for Hi-C datasets. Source data are provided with this paper.

## Code availability

Raw sequencing data generated on NextSeq550 or NextSeq2000 platforms were demultiplexed using TranslocPreprocess.pl from the HTGTS transloc pipeline, as described previously^43^. HTGTS-V(D)J-seq and 3C-HTGTS libraries were processed using the HTGTS transloc pipeline. 3C-HTGTS, GRO-seq and ChIP-seq datasets were further analyzed using the pipelines listed below. Hi-C datasets were processed with distiller-nf v0.3.4, and Hi-C scaling curves were generated using the pipeline listed below. Source code for all pipelines is available at the GitHub repositories listed below.

Transloc preprocessing and HTGTS-V(D)J-seq analysis: http://robinmeyers.github.io/transloc_pipeline/

3C-HTGTS analysis: https://github.com/Yyx2626/HTGTS_related/tree/main/3CHTGTS_related

GRO-seq analysis: https://github.com/Yyx2626/Fred_Alt_Lab/tree/master/GROseq

ChIP-seq analysis: https://github.com/Yyx2626/Fred_Alt_Lab/tree/master/ChIPseq

Hi-C processing: https://github.com/open2c/distiller-nf

Scaling curve processing: https://github.com/dekkerlab/Zhuo_Fred_Hi-C_SA12/blob/main

## Availability of materials

Plasmids and cell lines generated in this study are available from the laboratory of one of the corresponding authors (FWA) upon reasonable request.

## Extended Data Figure Legends

**Extended Data Fig. 1.**
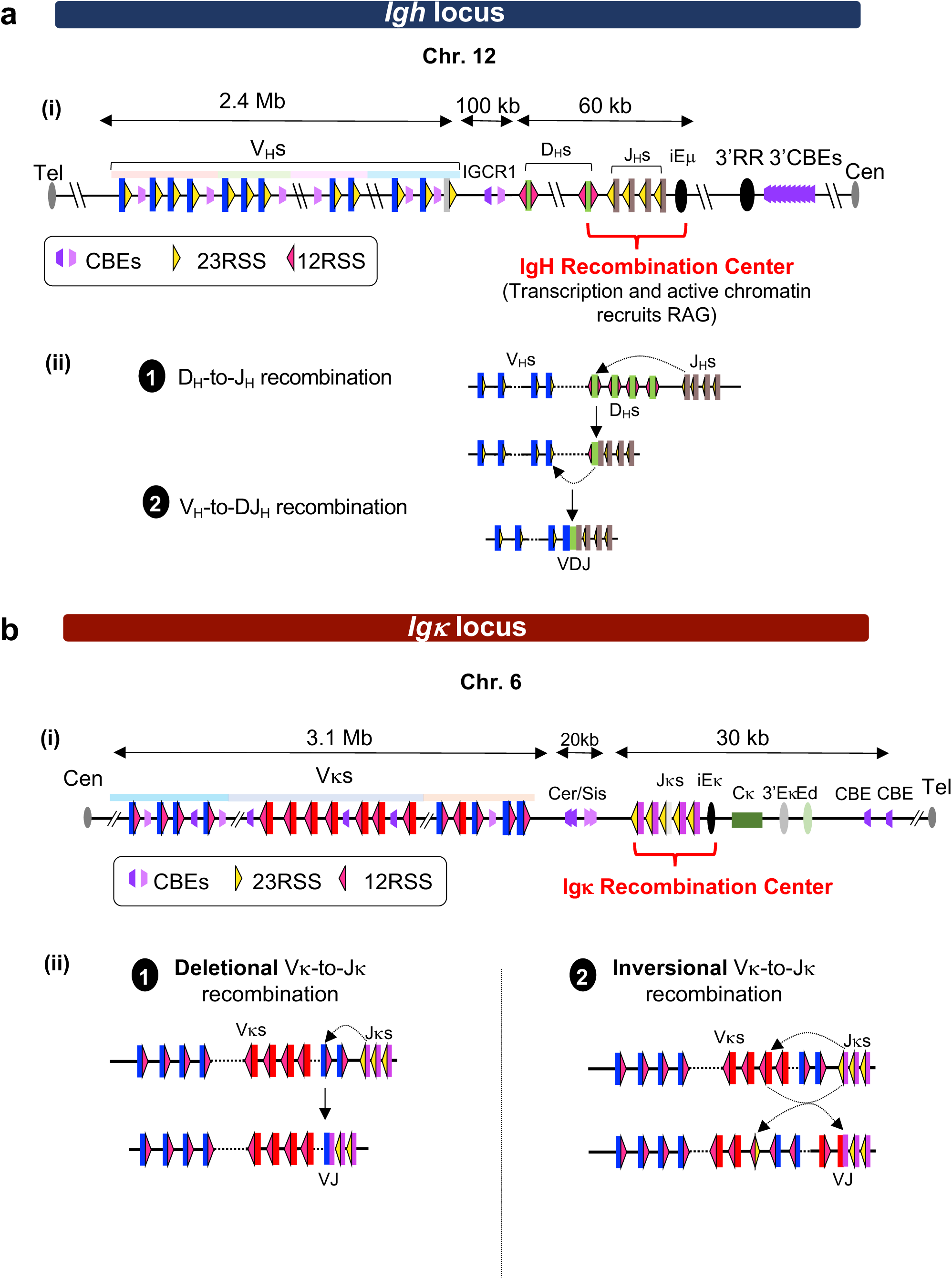
Organization and distinct recombination mechanisms of the *Igh* and *Igκ* loci. **a**, (**i**) Schematic of *Igh* (not to scale), comprising more than 100 V_H_s (blue box), 10 D_H_s (green box) and four J_H_s (brown box) segments across 2.7 Mb region. V_H_s families are distinguished by color. V_H_s and J_H_s segments are flanked by 23RSSs (yellow triangles), whereas D_H_ segments are flanked by 12RSSs (red triangles); triangle direction denotes RSS orientation. Trapezoids denote CBEs and their orientation. The J_H_ cluster and intronic enhancer form the *Igh* RC (red curly braces), and IGCR1 separates the V_H_ region from the DJ_H_ domain. Downstream 3’RR enhancers and upstream-oriented 3’CBEs are indicated. (**ii**) Sequential deletional D_H_-to-J_H_ and V_H_-to-DJ_H_ joining between convergently oriented 12RSS-23RSS pairs assembles a VDJ exon. **b**, (**i**) Schematic of *Igκ* (not to scale), containing more than 100 Vκs (blue and red box) segments across about 3 Mb and four functional Jκs (pink box) segments. Distal and proximal Vκ segments are predominantly oriented for deletional joining (blue), whereas middle Vκ segments are predominantly oriented for inversional joining (red). Vκs carry 12RSSs and Jκs carry 23RSSs. The Cer-Sis CBE platform lies upstream of the Jκ cluster and intronic enhancer iEκ, which form the primary *Igκ* RC (red curly braces). Further downstream, the Cκ, *Igκ* enhancers, upstream-oriented CBEs are indicated. (**ii**) Primary Vκ-to-Jκ joining can proceed by deletion or inversion.

**Extended Data Fig. 2.**
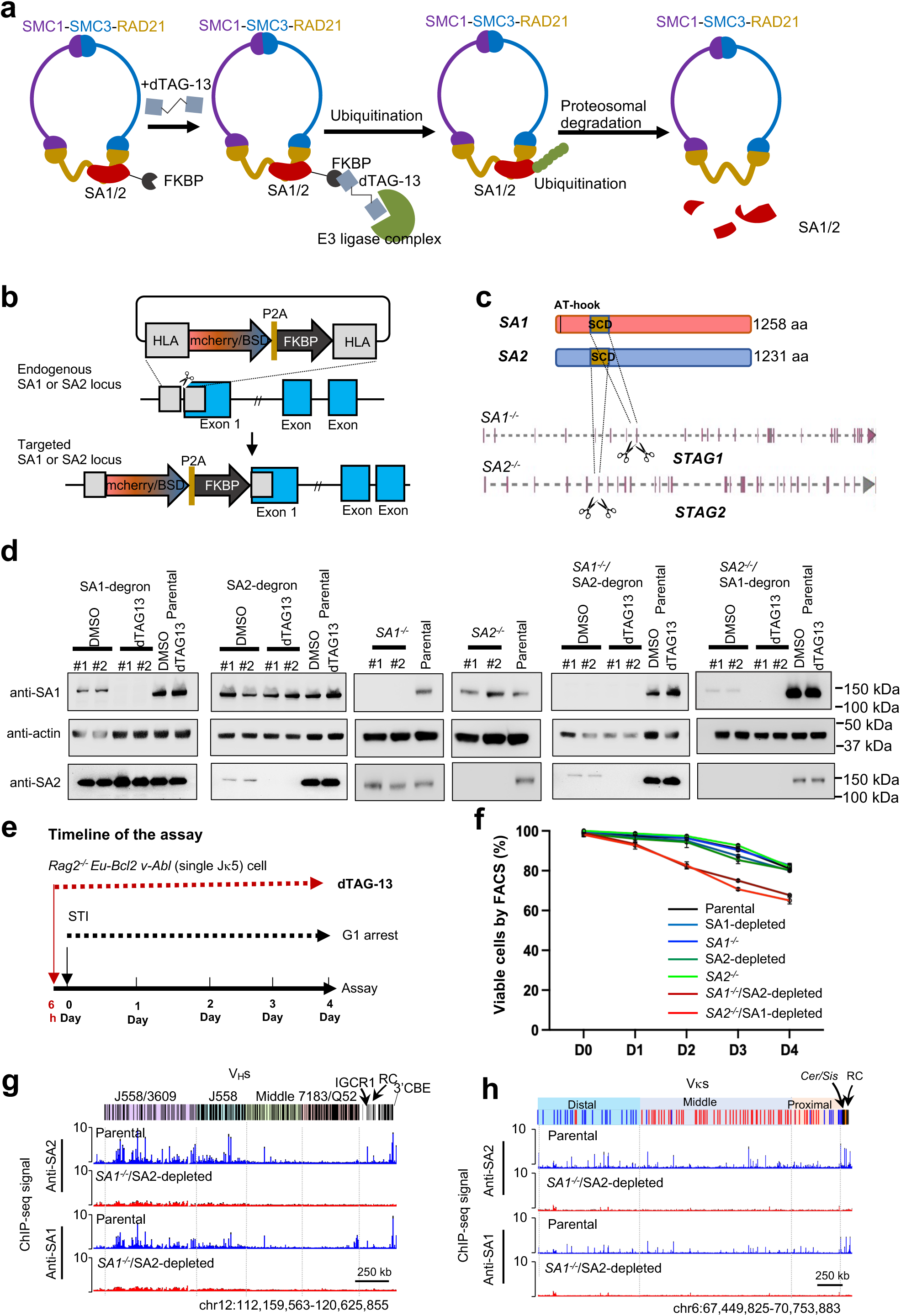
Generation and characterization of single and dual SA1 and/or SA2 inactivation cell lines. **a**, Schematic of dTAG-13-mediated degron depletion of SA1 and/or SA2. **b**, Strategy for engineering endogenous SA1 or SA2 alleles to encode N-terminal FKBP^F36V^ tags fused in-frame with either an mCherry reporter or blasticidin (BSD) resistance marker. The FKBP^F36V^ tag enables conditional degradation of tagged SA1 or SA2 upon dTAG-13 treatment, generating SA1-degron and SA2-degron lines. **c**, Schematic of SA1 or SA2 knockout by deletion of the Stromalin-Conserved Domain (SCD). SA1 and SA2 are encoded by *Stag1* and *Stag2* gene, respectively. The SCD is indicated by brown boxes, and corresponding exons are shown. Two sgRNAs used for SCD deletion are indicated by scissors. **d**, Western blot analysis of SA1 and/or SA2 protein levels in different genotypes of cell lines, compared with parental and DMSO-treated (untreated) controls. Antibodies are indicated on the left, and protein sizes are indicated on the right. **e**, Treatment scheme for dTAG-13 treatment and G0/G1 arrest of *v-Abl* cells. dTAG-13 was added 6 h before STI-571 treatment, and cells were then arrested in G0/G1 for 4 days. STI indicates STI-571. **f**, Cell viability analysis of the indicated genotypes during G0/G1 arrest. Cells were harvested daily over the 4-day arrest period and analyzed for viability. Viable-cell percentages at each time point and under each condition are shown as mean ± s.d. from three biologically independent samples. **g**,**h**, SA1 and SA2 ChIP-seq signals at the *Igh* (**g**) and *Igκ* (**h**) loci confirming efficient SA1/2 dual inactivation in *SA1^-/-^*/SA2-depleted cells. ChIP-seq signals are shown as mean ± s.e.m. from three biological replicates.

**Extended Data Fig. 3.**
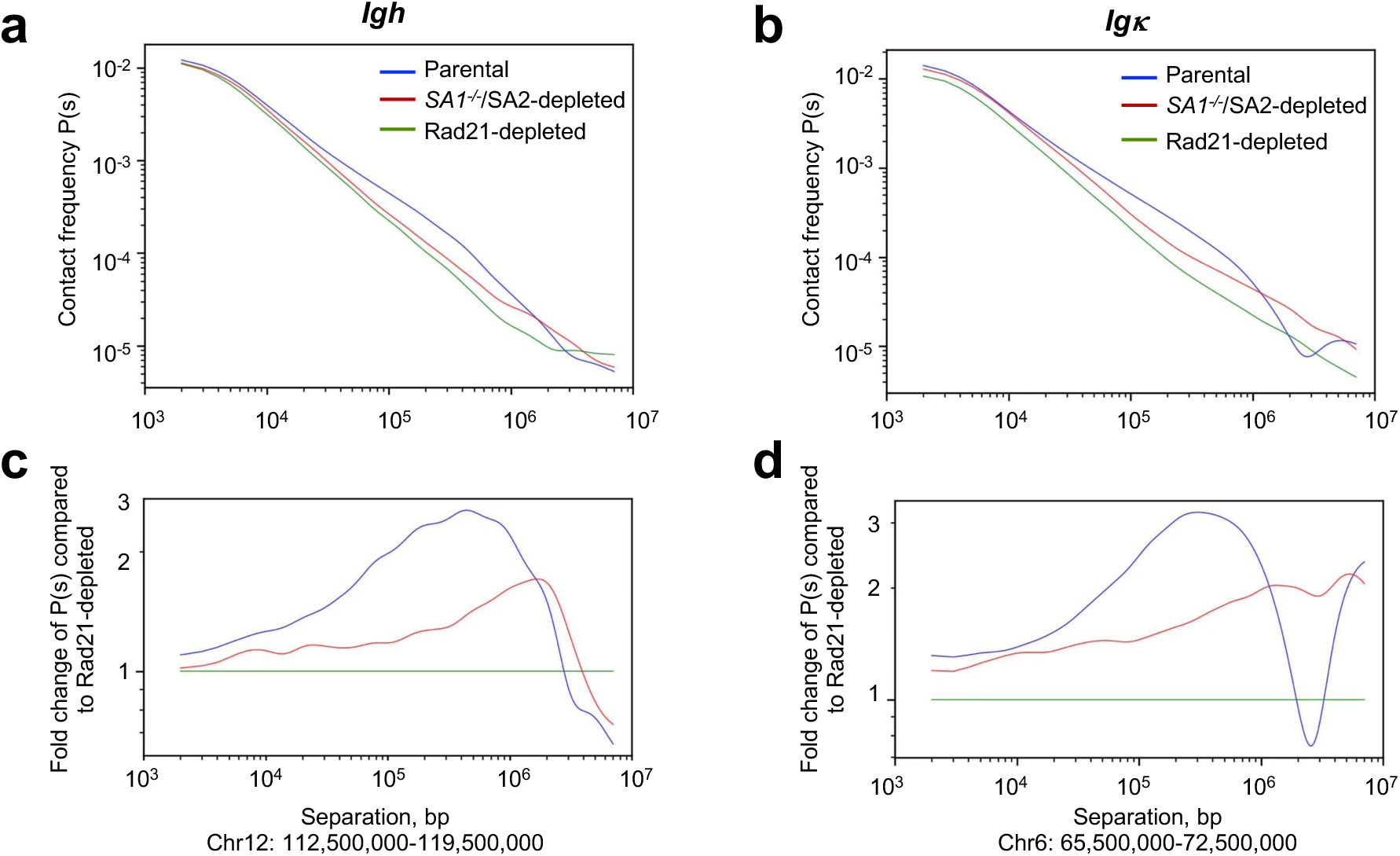
Contact probability decay analysis (P(s)) at the *Igh* and *Igκ* region. Related to Fig. 1**. a,b,** P(s) scaling plot of G0/G1-arrested cells at a region that include *Igh* locus (**a**) and the region that includes *Igκ* locus (**b**). Parental cell sample in blue line, *SA1^-/-^*/SA2-depleted sample in red line, and RAD21-depleted sample in green line. **c,d,** Fold change of P(s) shown in (**a** and **b**) comparing each condition with RAD21 depletion. chr., chromosome.

**Extended Data Fig. 4.**
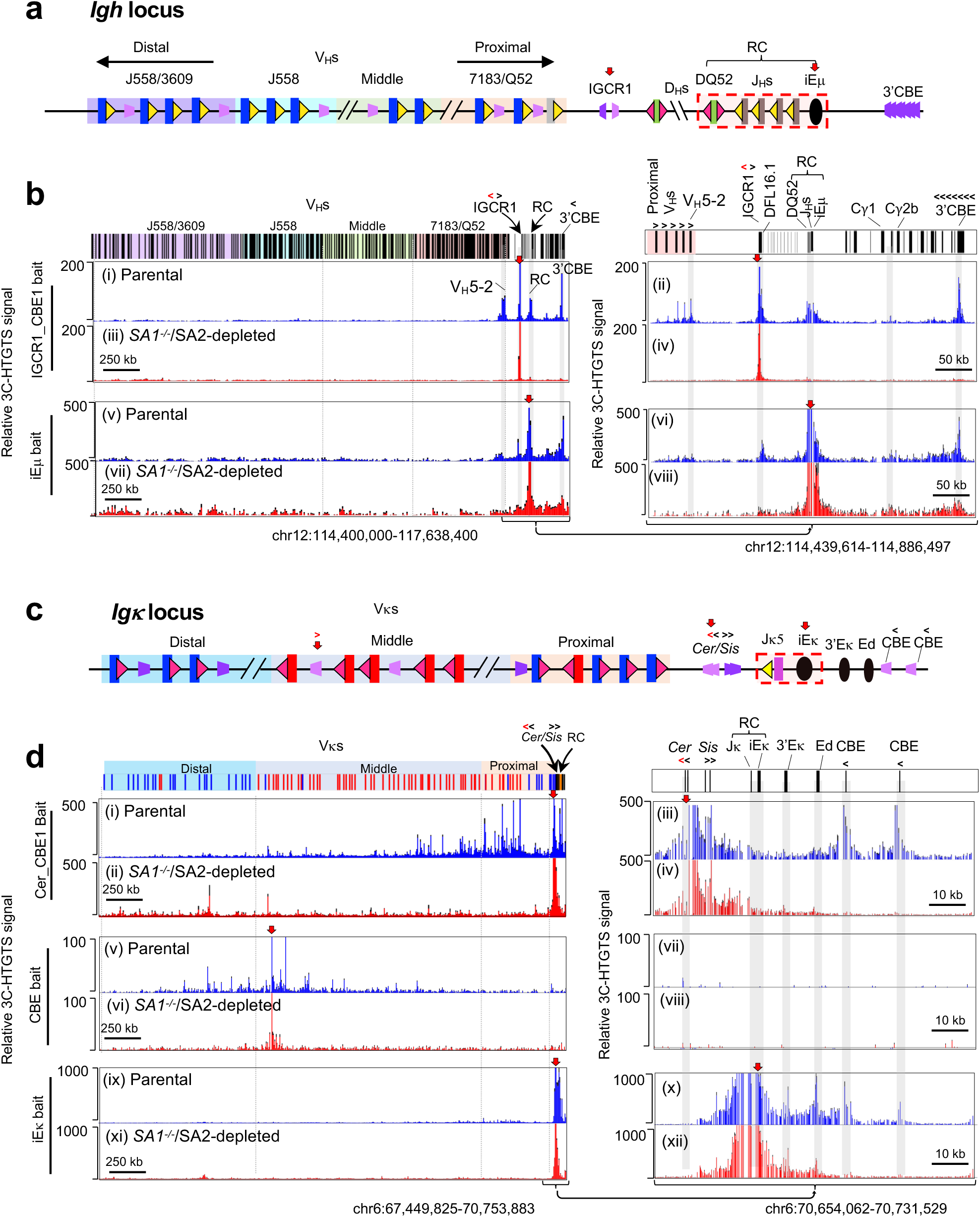
High-resolution 3C-HTGTS analysis of chromatin organization at the *Igh* and *Igκ* loci following SA1/2 inactivation. Related to Fig. 1. a, Schematic of the mouse *Igh* locus with the 3C-HTGTS bait labeled with red arrow at indicated IGCR1-CBE1 and iEμ sites. b, 3C-HTGTS interaction profiles across the four V_H_ domains and downstream region of the *Igh* locus using IGCR1-CBE1 (i-iv) and iEμ (v-viii) baits. The downstream *Igh* region is shown enlarged in the right panel. The orientation of CBEs is indicated with “<” point upstream and “>” point downstream. c, Schematic of the mouse *Igκ* locus, showing the 3C-HTGTS baits labeled with red arrow at Cer-CBE1, middle Vκ CBE, and iEκ sites. d, 3C-HTGTS interaction profiles at the *Igκ* locus using Cer-CBE1 (i-iv), middle Vκ CBE (v-viii), and iEκ (ix-xii) baits. The downstream *Igκ* region is shown enlarged in the right panel. Data in b,d are shown as mean ± s.e.m. from three biological replicates. chr., chromosome.

**Extended Data Fig. 5.**
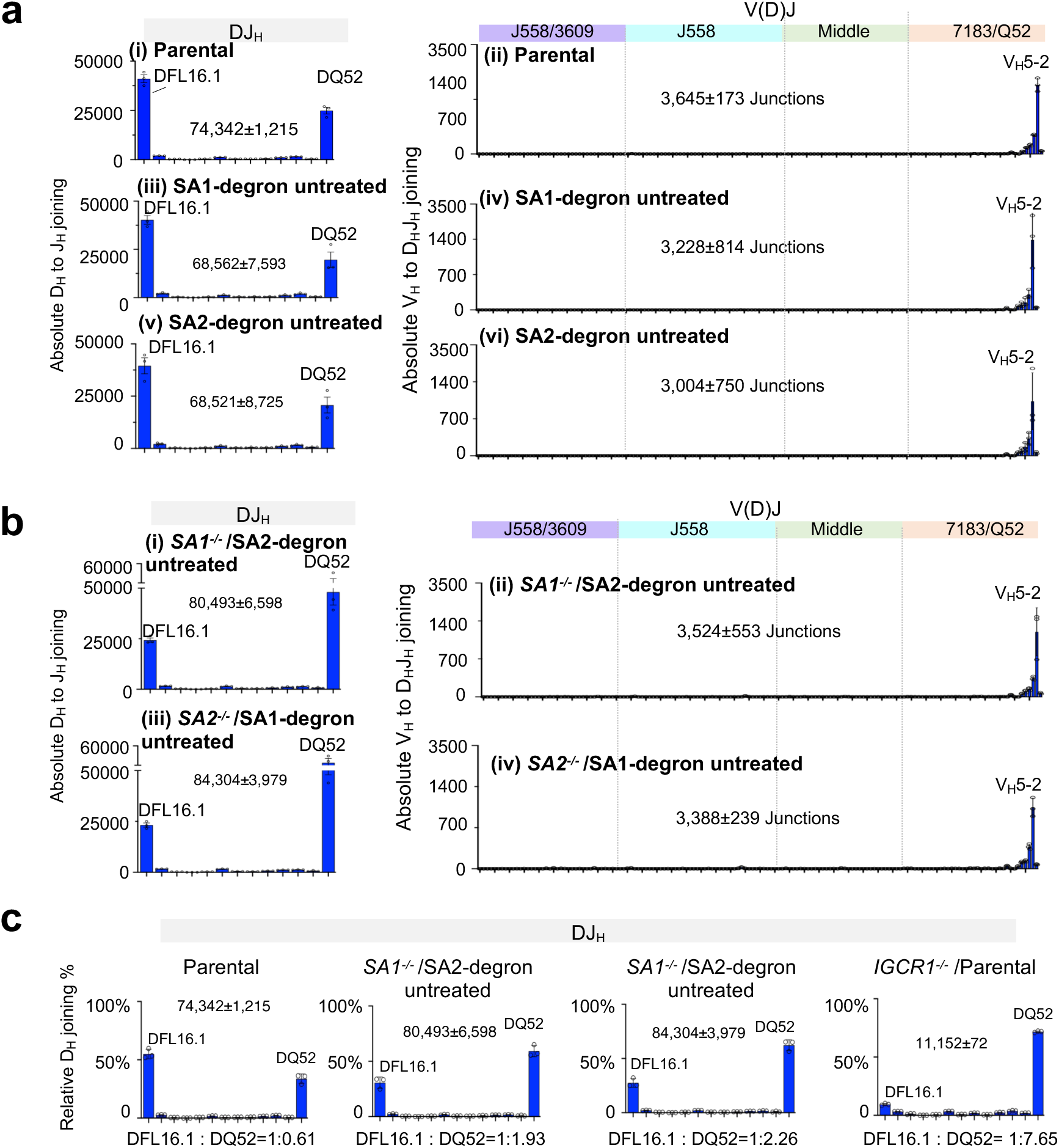
Impact of partial inactivation of SA1 and/or SA2 *v-Abl* cells in *Igh*. **a,b**, Absolute utilization of all *Igh* D_H_s (panels i,iii,v) and V_H_s (panels ii,iv,vi) in parental (**a.** i,ii) (reproduced from Fig. 2b for comparison), SA1-degron untreated cells (**a.** iii,iv), SA2-degron untreated cells (**a.** v,vi), *SA1^-/-^*/SA2-degron, and *SA2^-/-^*/SA1-degron untreated cells (**b**, i-iv). Deletional joins indicated in blue. Total junction numbers are shown in each panel and in subsequent figures. The overall rearrangements pattern of D_H_ and V_H_ in *Igh* are similar between parental and SA1-degron untreated or SA2-degron untreated cells. For D_H_ usage pattern compared with parental: r = 0.99 for SA1-degron untreated; r = 1.00 for SA2-degron untreated, with P < 0.001 for all comparisons. D_H_ usage patterns in untreated *SA1^-/-^*/SA2-degron and *SA2^-/-^*/SA1-degron cells differed modestly from parental cells, with Pearson correlation values of r = 0.81 and r = 0.78, respectively (P < 0.001 for both). For V_H_ usage pattern compared with parental: r = 1.00 for all indicated untreated lines, with P < 0.001. Moreover, D_H_ (r = 1.00, P < 0.001) and V_H_ (r = 0.84, P < 0.001) rearrangement patterns were same between *SA1^-/-^*/SA2-depleted and *SA2^-/-^*/SA1-depleted cells. There is no significantly different in total D_H_ rearrangement in parental with *SA1^-/-^*/SA2-degron or *SA2^-/-^*/SA1-degron untreated cells (P = 0.45; and P = 0.12, respectively). **c**, Relative D_H_ joining pattern of parental, *SA1^-/-^*/SA2-degron and *SA2^-/-^*/SA1-degron untreated, and *IGCR1^-/-^* cell line. DFL16.1:DQ52 ratios were calculated and are shown below each genotype panels. The corresponding P values are: parental, P = 0.0032; untreated *SA1^-/-^*/SA2-degron, P = 0.0027; untreated *SA2^-/-^*/SA1-degron, P < 0.001; and *IGCR1^-/-^*, P < 0.001. Locations of selected V_H_ segments are indicated. Data are presented as mean ± s.e.m. P values were calculated using an unpaired, two-tailed Welch’s t-test (n = 3).

**Extended Data Fig. 6.**
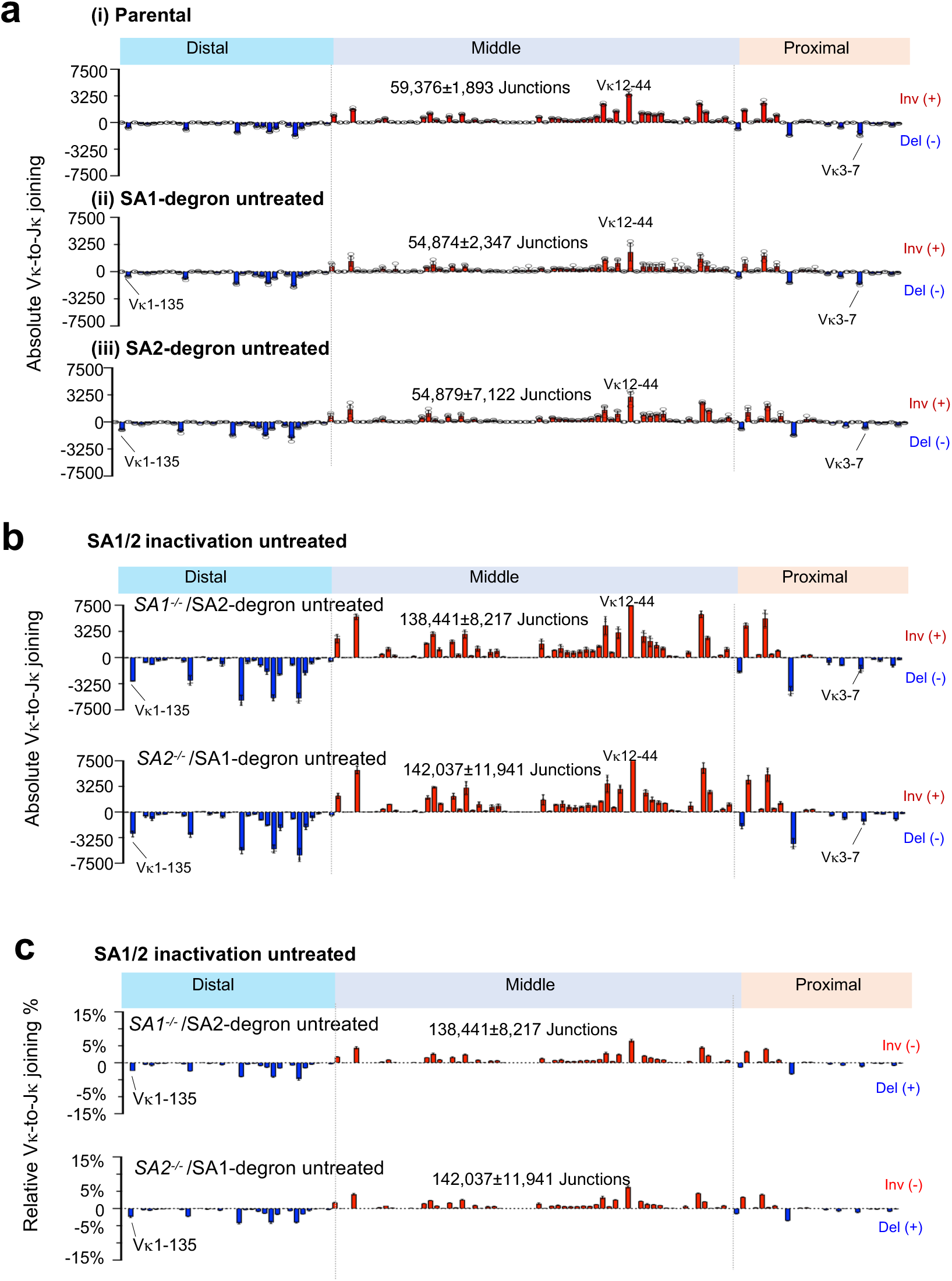
Impact of partial inactivation of SA1 and/or SA2 *v-Abl* cells in *Igκ*. **a,b**, Absolute utilization of all Vκs (panels i,ii,iii) in parental (**a.** i) (reproduced from Fig. 3b for comparison), SA1-degron untreated cells (**a.** ii), SA2-degron untreated cells (**a.** iii), *SA1^-/-^*/SA2-degron, and *SA2^-/-^*/SA1-degron untreated lines (**b**). Total junction numbers are shown in each panel. Total Vκ rearrangement pattern were similar between parental cells and untreated SA1-degron or SA2-degron cells, but the rearrangement increased more than two-fold in untreated SA1/2 dual-inactivation cells to parental cells: 2.33-fold in untreated *SA1^-/-^*/SA2-degron cells (P = 0.008) and 2.39-fold in untreated *SA2^-/-^*/SA1-degron cells (P = 0.018). Detailed values are provided in Supplementary Table 2. **c**, Relative Vκ joining pattern of SA1/2 dual inactivation untreated cell line. Statistical significance was determined using unpaired, two-tailed Welch’s t-tests (n = 3 biological replicates).

**Extended Data Fig. 7.**
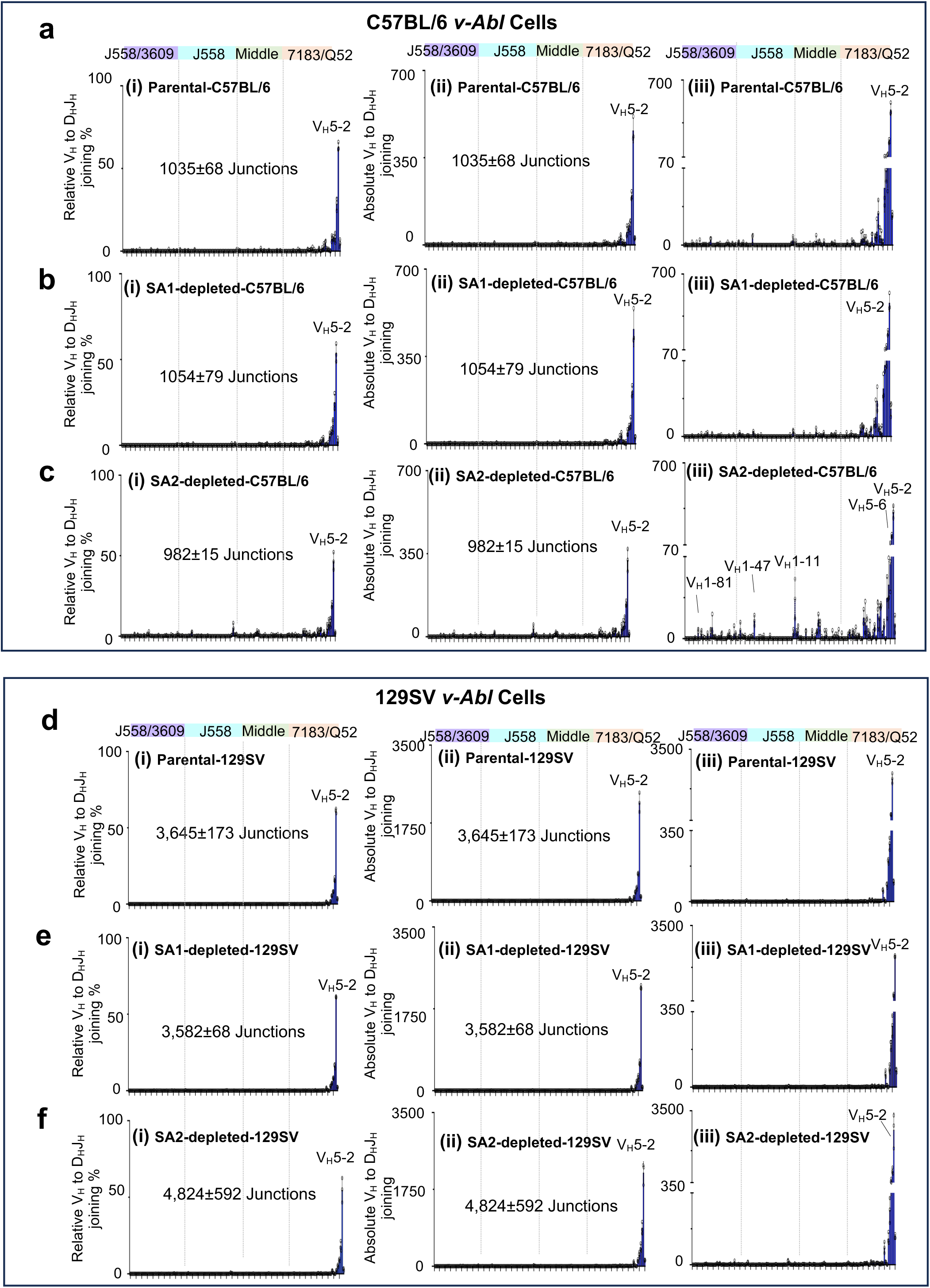
Impact of SA1 or SA2 inactivation on *Igh* V(D)J recombination in C57BL/6 *v-Abl* cells. Upper panels: C57BL/6-background *v-Abl* cells. **a-c**, V_H_-to-DJ_H_ joining in parental **(a),** SA1-depleted **(b)** and SA2-depleted **(c)** C57BL/6-background *v-Abl* cells. For each panel, subpanels i, ii, and iii show relative joining levels, absolute joining levels, and absolute joining levels plotted on a broken scale, respectively, to facilitate comparison with a recent study^40^ (see Methods for details). **Lower panels:** 129SV-background *v-Abl* cells. **d-f**, V_H_-to-DJ_H_ joining in parental **(d**), SA1-depleted **(e),** and SA2-depleted **(f)** 129SV-background *v-Abl* cells. Subpanels in panel **d-f** are organized as described for panel **a-c**. In parental C57BL/6-background cells, the ratio of V_H_5-2 usage to the combined usage of the 12 most proximal V_H_s immediately distal to V_H_5-2 was (0.95 ± 0.04):1, compared with (1.81 ± 0.05):1 in parental 129SV-background cells (**a,d**, subpanel i), confirming more dominant V_H_5-2 usage in the 129SV background (P = 0.0003). The ratio of V_H_5-2 usage to that of the total upstream V_H_s is (0.80 ± 0.03):1 in parental C57BL/6-background cells and (0.49 ± 0.05):1 in SA2-depleted C57BL/6-background cells, which confirms a low, but significant (P = 0.0092) increase in upstream V_H_ rearrangement upon SA2-inactivation in C57BL/6-background cells (**a,c**, subpanel i). There was is no significant difference in ratios of V_H_5-2 usage to upstream V_H_ rearrangements versus that of parental cells ((1.57 ± 0.06):1) in SA1-depleted ((1.58 ± 0.02):1, P = 0.8847), and SA2-depleted ((1.56 ± 0.08):1, P = 0.9053) 129SV-background *v-Abl* cells (**d-f**, subpanel i). Data are presented as mean ± s.e.m. Statistical significance was determined using unpaired, two-tailed Welch’s t-tests (n = 3 biological replicates).

**Extended Data Fig. 8.**
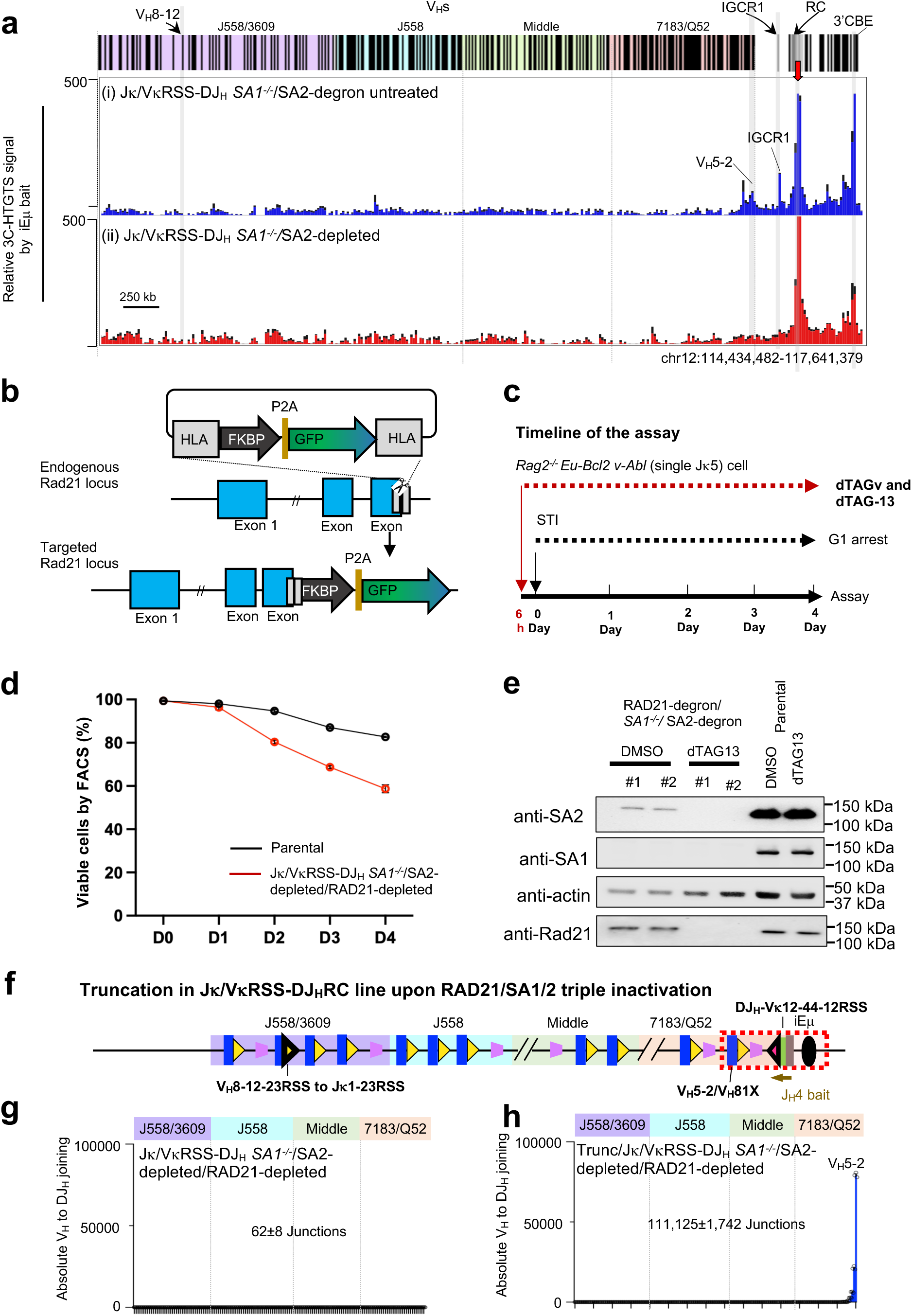
*Igκ* RSS promoted *Igh* V_H_-to-DJ_H_ joining requires loop extrusion. Related to Fig. 5**. a,** 3C-HTGTS interaction profiles using iEμ bait in double *Igκ* RSS replacement in Jκ/VκRSS-DJ_H_ *SA1^-/-^*/SA2-degron cell lines with or without dTAG-13 treatment. Red arrows indicate bait locations. chr., chromosome. 3C-HTGTS data are presented as mean ± s.e.m. of three biological replicates. **b,** Strategy for engineering endogenous RAD21 alleles to encode C-terminal FKBP^F36V^ tags fused in-frame with GFP reporter. **c**, Treatment scheme for combined dTAG-13 and dTAGv treatment in G0/G1 arrest of *v-Abl* cells. STI indicates STI-571. **d**, Cell viability analysis. Treatment scheme for combined dTAG-13 and dTAGv treatment 6 hours before STI G0/G1 arrest in *v-Abl* cells. **e**, Western blotting analysis of RAD21, SA1, and SA2 protein levels in Jκ/VκRSS-DJ_H_ *SA1^-/-^*/SA2-degron/RAD21-degron triple inactivation cell lines after 6 h dTAG-13 plus dTAGv treatment. Antibodies are indicated on the left, and protein sizes are indicated on the right. **f**, Diagram illustrating the truncation the 100 kb interval between V_H_5-2 and the DJ_H_ RC to generate the Trunc/Jκ/VκRSS-DJ_H_ *SA1^-/-^*/SA2-degron/RAD21-degron *v-Abl* cell line. **g**,**h**, Absolute utilization of V_H_s in *Igh* under Jκ/VκRSS-DJ_H_ *SA1^-/-^*/SA2-deplted/RAD21-depleted (**g**) and Trunc/Jκ/VκRSS-DJ_H_ *SA1^-/-^*/SA2-deplted/RAD21-depleted (**h**) *v-Abl* cell lines. Junction numbers are shown in each. Locations of selected V_H_ segments are indicated. V_H_ utilization is shown as mean ± s.e.m. from three biological replicates.

**Extended Data Fig. 9.**
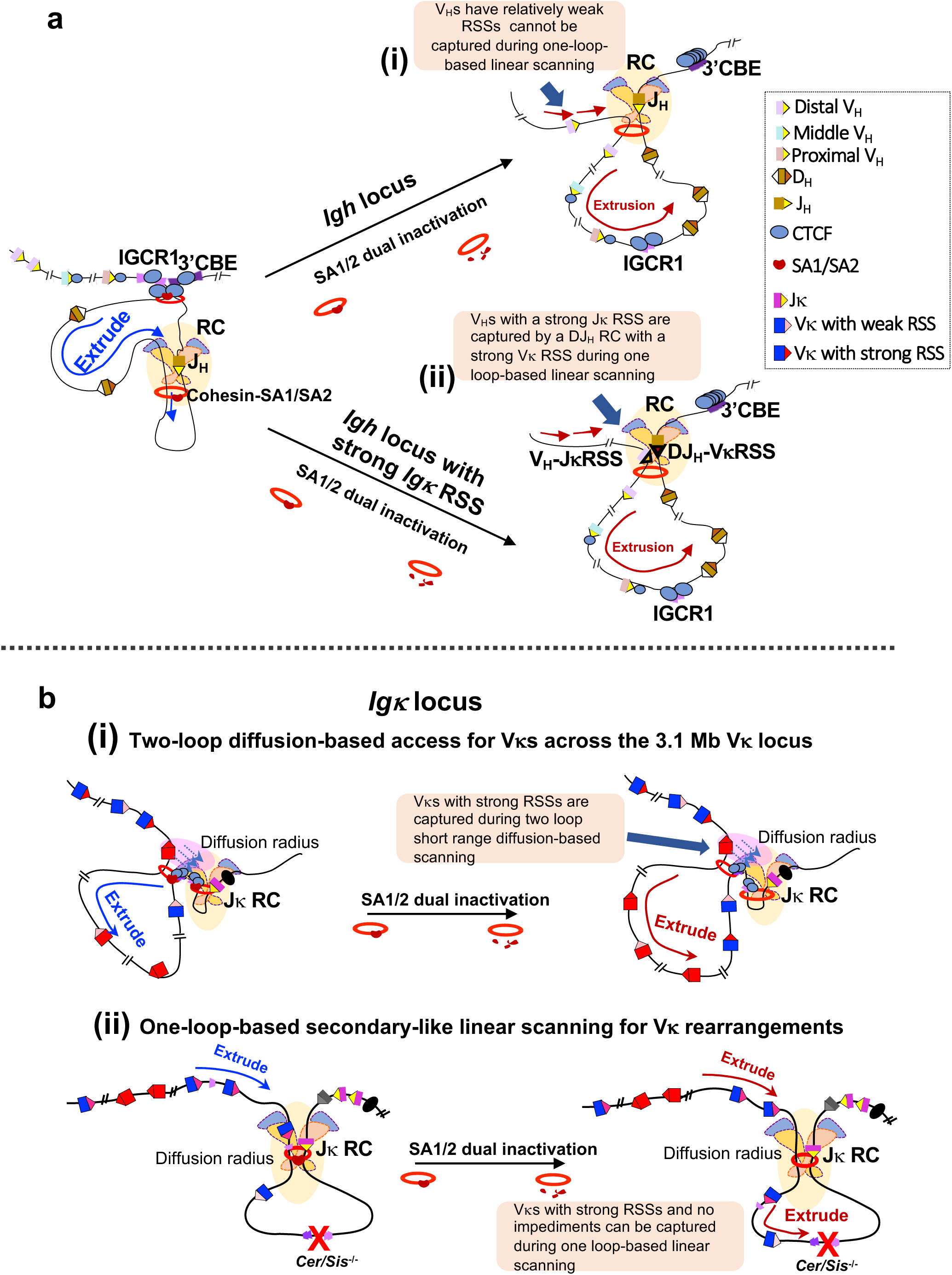
Working model. **a**, (i) Weak *Igh* RSSs require associated loop extrusion impediments, to increase their interactions with RC and capture for V(D)J recombination. These impediments are neutralized by SA1/2 dual inactivation, which abrogates linear-scanning mediated *Igh* V(D)J recombination. (ii) The very strong *Igκ* RSSs allow a V_H_ to be captured by the RAG-bound, J_H_-based RC during linear RAG scanning in the context of loop-extrusion impediment neutralization by SA1/2 dual inactivation. **b,** (i) The Cer/Sis diffusion platform mediates two-loop-based based primary diffusional Vκ to Jκ joining; (ii) Deletion or displacement of Cer/Sis allows one-loop based secondary-like linear scanning. Most primary (i) and secondary (ii) Vκ to Jκ rearrangements are not affected by absence of scanning impediments upon their neutralization by SA1/2 dual inactivation.

