## Supplementary Information.zip for "Robust V(D)J Recombination in the Absence of SA1/2-mediated Chromatin Loops": Supplementary information 1.pdf

**Supplementary information Fig. 1 | Features of the impediments neutralized by SA1/2 dual inactivation. Related to Fig. 1 and Extended data Fig.4. a-d,** Colocalization analysis of 3C-HTGTS interaction peaks with transcription, cohesin-RAD21, and CTCF. Black dashed lines indicate Cer-CBE1-baited interaction peaks that colocalize with transcription, cohesin-RAD21, and/or CTCF. Yellow dashed lines indicate middle Vκ CBE-baited interaction peaks that colocalize with transcription, cohesin-RAD21 and/or CTCF. Genomic features underlying each indicated peak are annotated as CBE (C), E2A-binding site (E), or transcription-associated site (T). Signals in **a, b** and **d** from parental cells are shown in blue, and signals from *SA1*$^{-/-}$/SA2-depleted cells are shown in red. **a,b**, 3C-HTGTS interaction profiles across the Vκ locus using Cer CBE1 (**a**) or a middle Vκ CBE (**b**) as bait in RAG-deficient parental and *SA1*$^{-/-}$/SA2-depleted *v-Abl* cells. Cer CBE1 and middle Vκ CBE-associated interaction peaks and their underlying features across the Vκ locus are indicated. **c**, GRO-seq data in RAG-deficient parental and *SA1*$^{-/-}$/SA2-depleted *v-Abl* cells. Signals on the positive and negative DNA strands are shown in red and blue, respectively. **d**, RAD21 and CTCF ChIP-seq signal in RAG-deficient parental (blue) and *SA1*$^{-/-}$/SA2-depleted (red) *v-Abl* cells. E2A binding was annotated based on published E2A ChIP-seq data from RAG-deficient primary pro-B cells. 3C-HTGTS, Gro-seq, and ChIP-seq data are presented as mean ± s.e.m. of three biological replicates.

### Supplementary information Fig.1

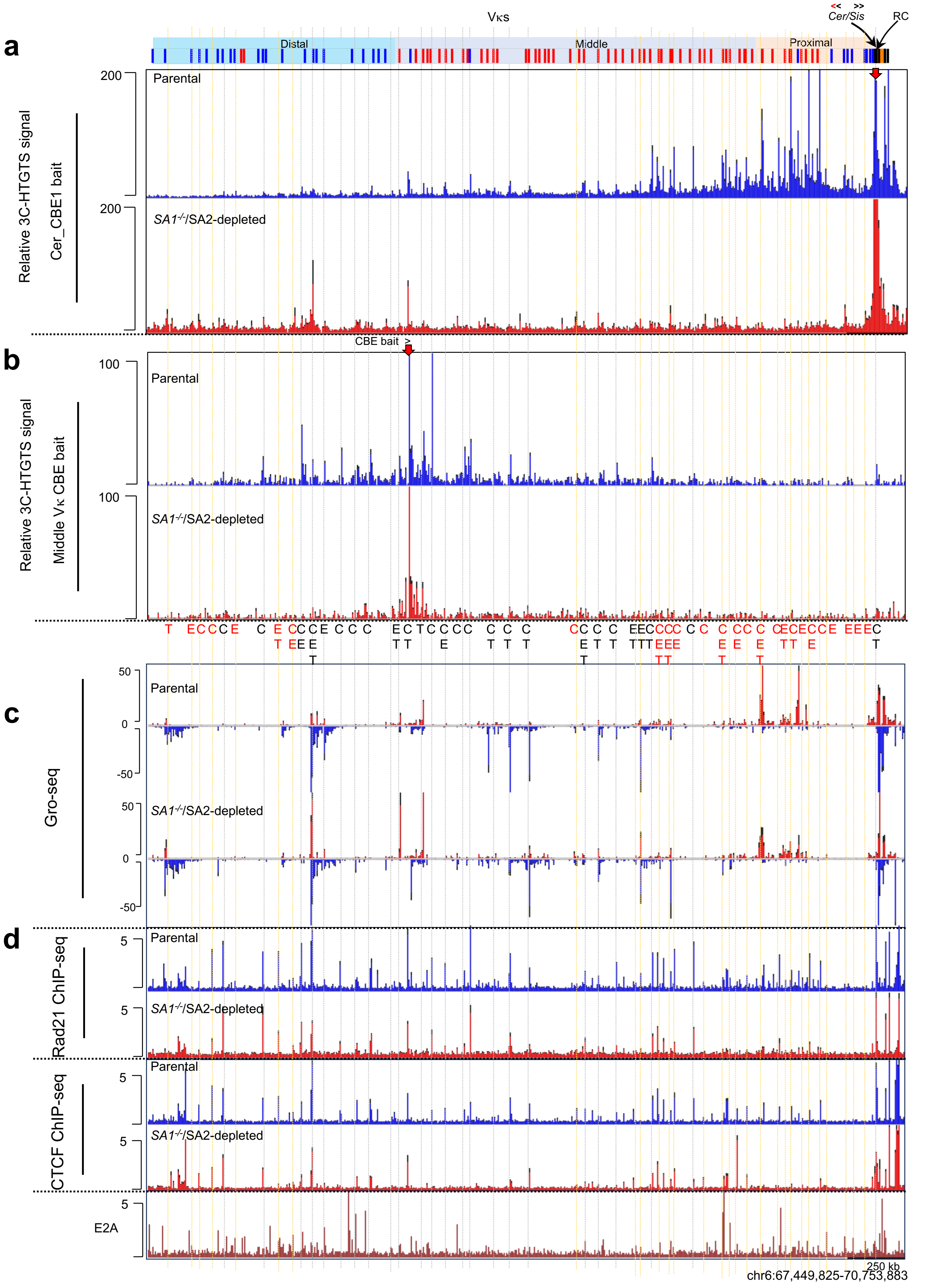
