## Supplementary Information.zip for "Robust V(D)J Recombination in the Absence of SA1/2-mediated Chromatin Loops": Supplementary information 2.pdf

**Supplementary information Fig.2 | RAD21-depletion abrogates *Igκ* Cer CBE1 interaction, RC transcription and *Igκ* V(D)J rearrangement. a**, Schematic of the mouse *Igκ* locus, showing the 3C-HTGTS bait labeled with red arrow at Cer CBE1. **b**, 3C-HTGTS interaction profiles across the Vκ region (i-ii) and downstream region (iii-iv) of the *Igκ* locus using Cer CBE1 baits upon RAD21-degron untreated and depletion treated *v-Abl* cells. The downstream *Igκ* region is shown enlarged in the below panel (iii-iv). The orientation of CBEs is indicated with "<" point upstream and ">" point downstream. **c**, GRO-seq data in RAG-deficient RAD21-degron untreated, RAD21-depleted, parental, and *SA1*$^{-/-}$/SA2-depleted *v-Abl* cells. Signals on the positive and negative DNA strands are shown in red and blue, respectively. **d**, Absolute utilization of all Vκs in *Igκ* locus in RAD21-degron untreated and depletion treated *v-Abl* cells. Deletional Vκ-Jκ1 joins are plotted below the x-axis in blue and labeled "Del (−)", inversional Vκ-Jκ1 joins are plotted above the x-axis in red and labeled "Inv (+)". Total junction numbers are shown in each panel. Data are shown as mean ± s.e.m. from three biological replicates. chr., chromosome.

**Further Discussion of Supplementary information Fig.2.**

In contrast to SA1/2 dual inactivation, RAD21 inactivation abrogated *Igκ* RC transcription that is required for its function[28] (panel c). RC activity depends on activity of three enhancers[53]: the iEκ located in the RC[54], the 3'Eκ enhancer 10 kb downstream[55, 56], and potentially the Ed enhancer 20 kb downstream[57]. While the mechanism by which RAD21 inactivation abrogates *Igκ* RC transcriptional activity is unknown, it is likely that the full *Igκ* RC enhancer complex is assembled by loop extrusion potentially in association with downstream *Igκ* downstream CBE approximately 30 kb downstream[53] and, therefore, inactivated by the absence of loop extrusion in the RAD21-inactivated background. We note that the iEμ-based *Igh* RC does not require distal elements brought together for loop extrusion and retains activity in the in the RAD21-depleted background[13].

### Supplementary information Fig. 2

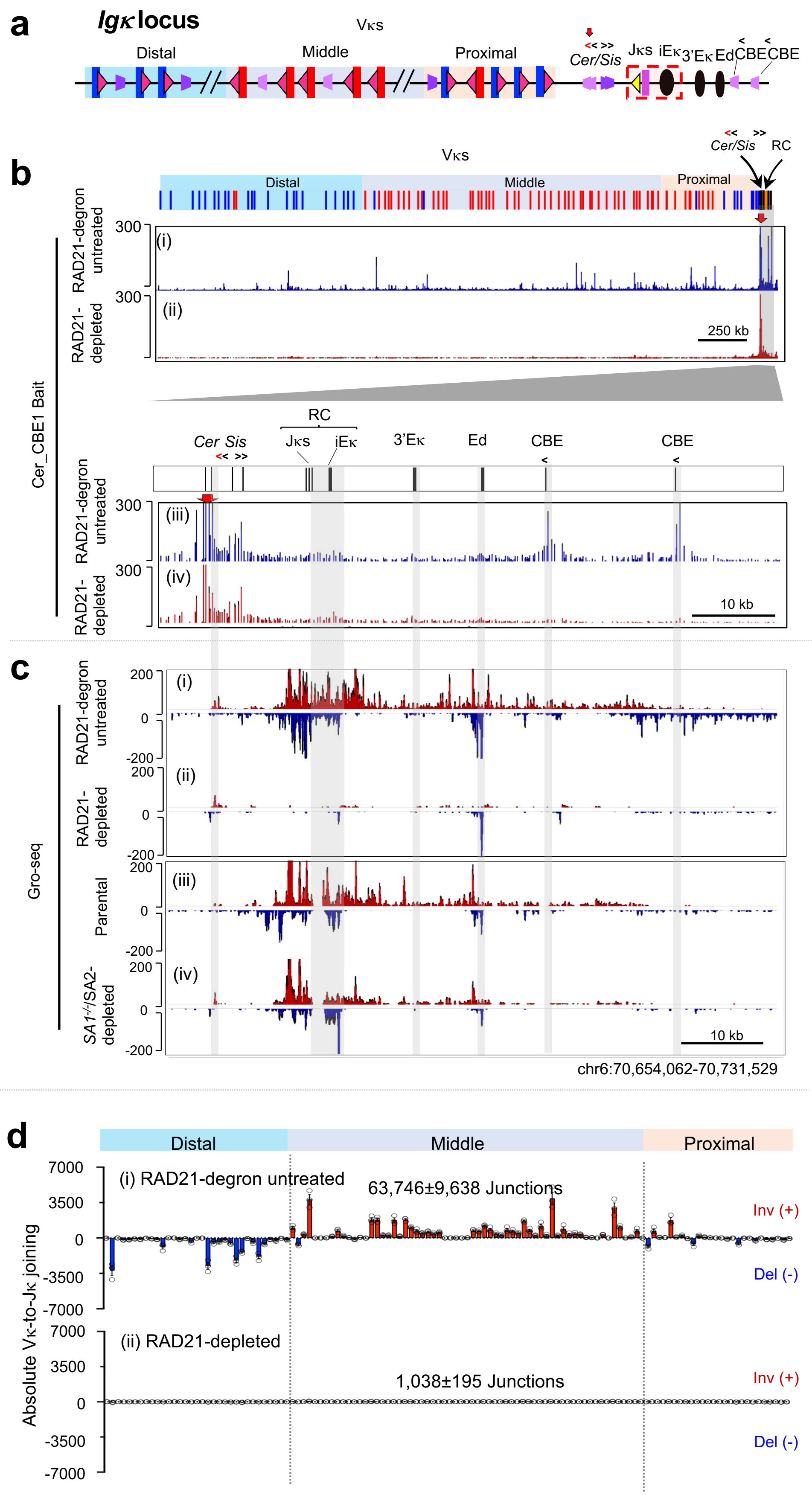
