## Supplementary Information.zip for "Robust V(D)J Recombination in the Absence of SA1/2-mediated Chromatin Loops": Supplementary information 3.pdf

**Supplementary information Fig.3 | Relative Vκ rearrangement pattern in single and double SA1 and/or SA2 inactivation. Related to Fig. 3. a-d, panel i,** Relative Vκ to Jκ rearrangement in parental (**a**), *SA1*$^{-/-}$ (**b**; upper panel), SA1-depleted (**b**; lower panel), *SA2*$^{-/-}$ (**c**; upper panel), SA2-depleted (**c**; lower panel), *SA1*$^{-/-}$/SA2-depleted (**d**; upper panel) and *SA2*$^{-/-}$/SA1-depleted (**d**; lower panel) *v-Abl* cells. Junction counts are indicated in each panel and in subsequent figures as to allow comparison to absolute rearrangement levels. **a-d, panel ii,** Utilizations percentage of distal Vκ "D" in blue bar, middle Vκ "M" in gray bar, and proximal Vκ "P" in pink bar. Notably, SA1/2 dual inactivation significantly shifted Vκ rearrangement pattern relative to parental cells, decreasing proximal Vκ usage 2.12-fold for *SA1*$^{-/-}$/SA2-depleted and 1.76-fold for *SA2*$^{-/-}$/SA1-depleted (both $P < 0.001$), and increasing distal Vκ usage 2.16-fold for *SA1*$^{-/-}$/SA2-depleted and 2.08-fold for *SA2*$^{-/-}$/SA1-depleted (both $P < 0.001$). Statistical significance was determined using unpaired, two-tailed Welch's t-tests (n = 3 biological replicates): *$P \leq 0.05$, **$P \leq 0.01$ and ***$P \leq 0.001$.

### Supplementary information Fig.3

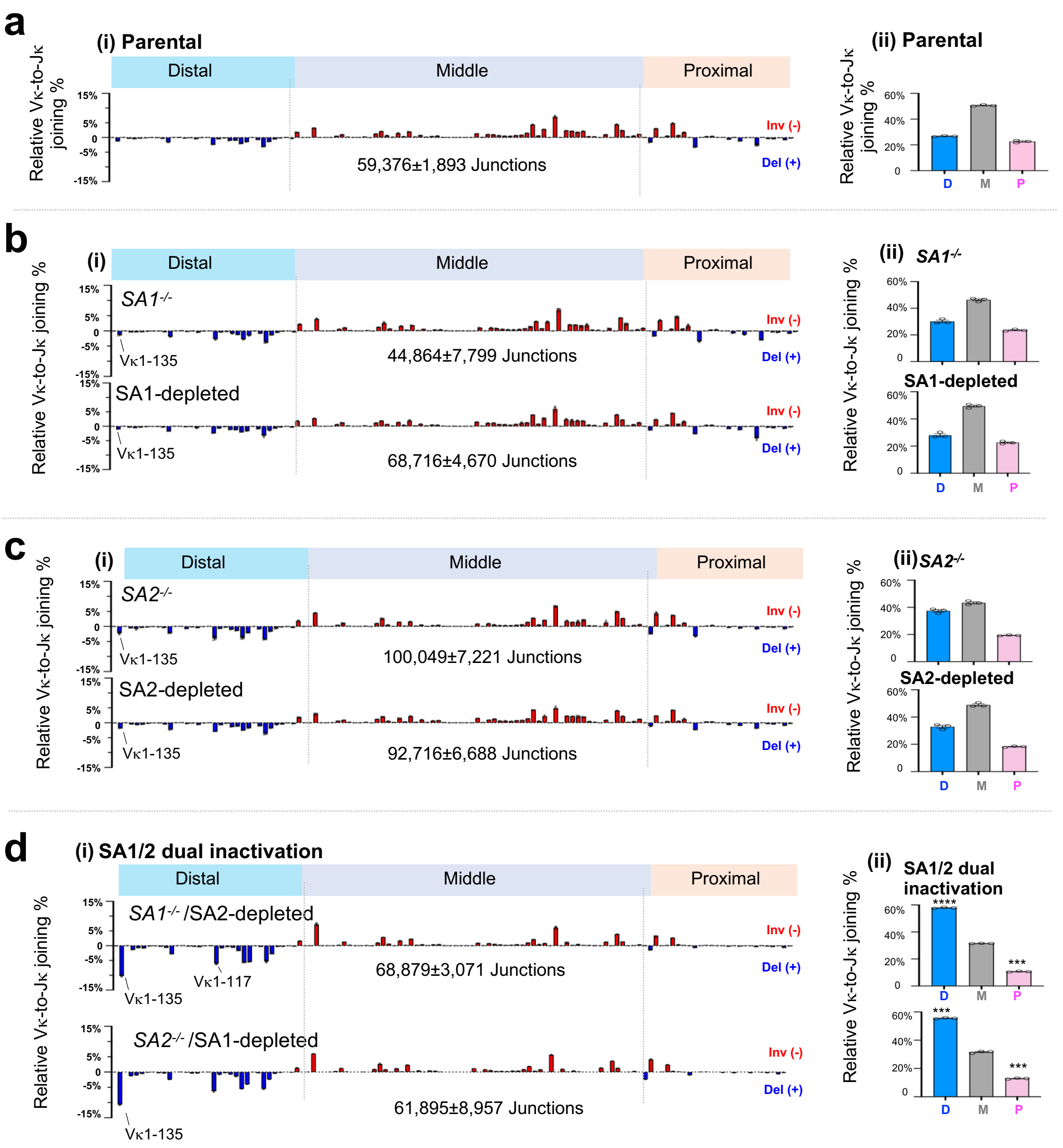
