## Supplementary Information.zip for "Robust V(D)J Recombination in the Absence of SA1/2-mediated Chromatin Loops": Supplementary information 4.pdf

**Supplementary Figure 4 | TdT gene expression and protein level in 129SV-background *v-Abl* cells.** **a**, TdT gene (*dntt*) expression level shown in Gro-seq data in G0/G1-arrested bone marrow pro-B cell, G0/G1-arrested 129SV-*v-Abl* parental cells, and G0/G1-arrested 129SV-*v-Abl* $SA1^{-/-}$/SA2-depleted cells. **b**, Western blotting analysis of TdT protein level in 129SV-background *v-Abl* cell lines: parental, SA1-depleted, SA2-depleted, and $SA1^{-/-}$/SA2-depleted cells. Total thymocytes cells were used as positive TdT protein control. Antibodies are indicated on the left, and protein sizes are indicated on the right.

### Supplementary Figure 4

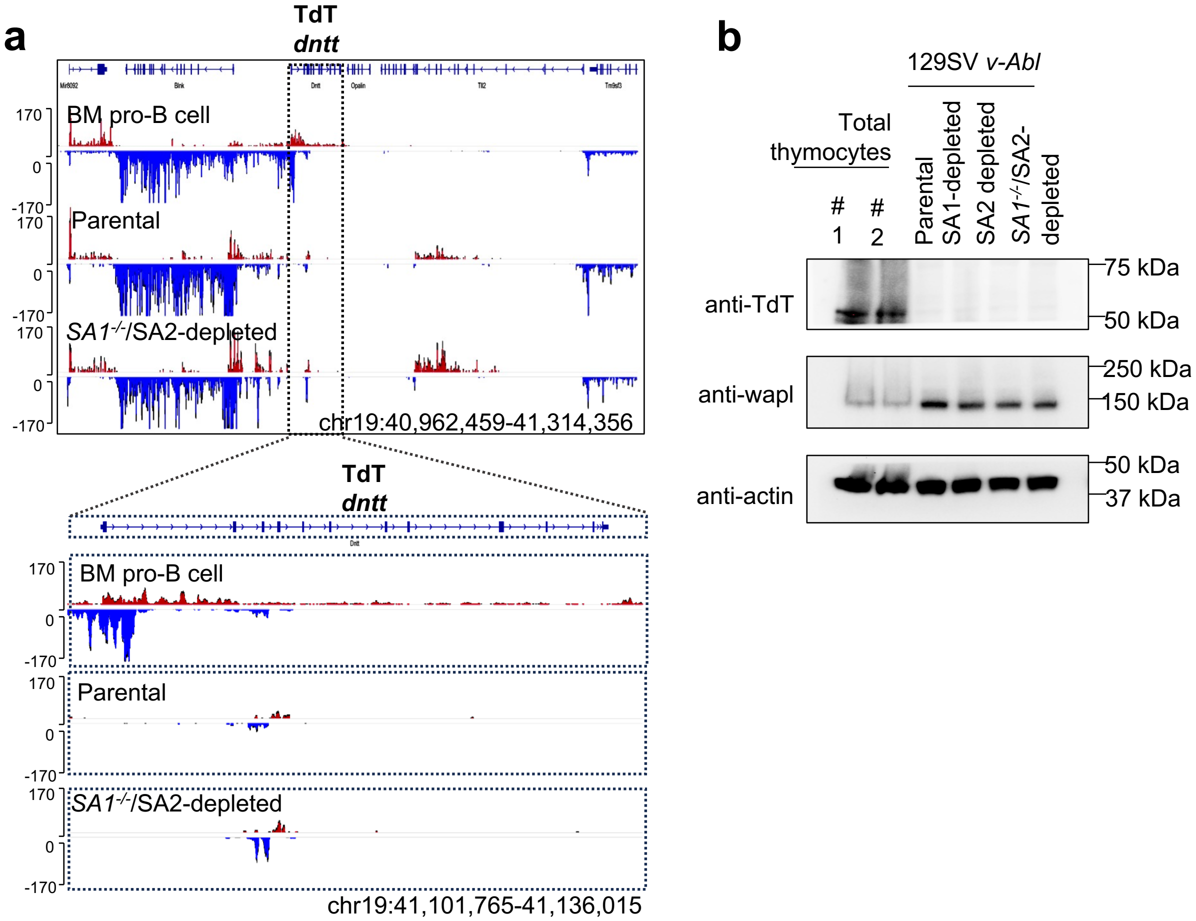
